# Partial input loss differentially modifies neural pathways

**DOI:** 10.1101/2025.02.14.638366

**Authors:** Joo Yeun Lee, Jeffrey Yang, Scott C. Harris, Jeremiah V. John, Natalia S. Stone, Luca Della Santina, David B. Kastner, Felice A. Dunn

**Author notes:** Corresponding authors contacts.

## Abstract

Following input loss from degeneration, injury, and/or aging, downstream circuits undergo modifications that can impact neural computations. How neural computations across different pathways are affected by common input loss remain understudied. To leverage known cell types, well-defined circuitry, and molecular tools, we use the mouse retina to show how multiple pathways adjust their functional properties differently to common input loss and further locate these changes within each pathway. Specifically, we asked if two OFF ganglion cell types, alpha OFF-sustained (A_OFF-S_) and OFF-transient (A_OFF-T_) cells, and their respective dominant presynaptic partners, type 2 and type 3a cone bipolar cells, respond differentially to partial cone loss. We find that A_OFF-T_ ganglion cells exhibit more circuit changes than A_OFF-S_ ganglion cells, resulting in altered spatiotemporal tuning following partial cone loss. We show that the underlying mechanisms include changes in glutamatergic, GABAergic, and glycinergic circuits in the pathway of A_OFF-T_ ganglion cells. In response to common input loss, our study finds different locations of circuit modifications across OFF pathways. These findings provide insight into how sensory pathways can compensate differentially to common input loss.

## INTRODUCTION

Across sensory systems, neuronal pathways can undergo distinct changes in response to deafferentation (Villa et al., 2016; Liu et al., 2017; Eavri et al., 2018; Karoui et al., 2023; Kumar et al., 2023). The mechanisms underlying these differential responses and the implications for potential pathway-specific repair remain unknown. Given the dynamic properties of our nervous system even when natural regeneration is not possible, achieving optimal sensory restoration through intervention requires systematic examination of the states of remaining circuitry in response to deafferentation. Our understanding of that degeneration process is severely limited: while we know that neurons can die, we lack understanding of how the death of neurons in one part of a circuit influences processing downstream. Using the retina’s well-characterized circuits, we precisely controlled the degree of cone death to understand whether partial cone loss has pathway-specific effects downstream. Visual signals transmitted from photoreceptors diverge at the dendrites of bipolar cells, the second-order neurons of the mammalian retina, into 11-15 pathways (Shekhar et al., 2016; Peng et al., 2019; Hahn et al., 2023). These secondary pathways further diverge into 17 to >30 types of ganglion cells with diverse functional properties (Chander and Chichilnisky, 2001; Kim and Rieke, 2001; Chichilnisky and Kalmar, 2002; Murphy and Rieke, 2006; Puthussery et al., 2009; Nirenberg et al., 2010; Baden et al., 2016; Grünert and Martin, 2021; Goetz et al., 2022). In injury and disease, specifically animal models of optic nerve crush, glaucoma, and retinitis pigmentosa, certain ganglion cell types tend to be more vulnerable (O’Brien et al., 2014; Ou et al., 2016; Tran et al., 2019a; Wang et al., 2021; Dyszkant et al., 2025a), demonstrating differential effects across cell types. However, how different ganglion cell types differentially respond to common input loss, and how the circuit physiology and connectivity change before ganglion cell degeneration remains unknown. Answering this question has important implications for understanding how downstream neurons respond to common input loss, which occurs in normal aging, neurodegeneration, and injury. Therefore, we asked how different pathways change upon photoreceptor loss. Previously, we demonstrated how the alpha ON-sustained (A_ON-S_) ganglion cells react to partial loss of cones in the mature retina (Care et al., 2019; Lee et al., 2022). The loss of cones caused changes to the spatial receptive field of A_ON-S_ ganglion cells, including narrower centers and wider surrounds (Care et al., 2019; Lee et al., 2022). In this study, we ask if two OFF pathways exhibit similar characteristics or undergo unique pathway-specific changes after cone loss. We find evidence for modifications across retinal circuits, e.g., differential changes in excitatory and inhibitory inputs to two A_OFF_ ganglion cells, suggesting that the retina undergoes pathway-specific changes following partial cone loss. Such results are significant in demonstrating differential reactions to common input loss, their underlying mechanisms, and the potential for rescue across multiple pathways for sensory processing.

## RESULTS

### Partial cone loss causes differential temporal changes to A_OFF_ ganglion cell types

To determine if two OFF pathways exhibit global light-adapted characteristics or undergo unique pathway-specific changes after cone loss, we examined the anatomical and functional properties across retinal pathways. We focus on two types of OFF ganglion cells: the alpha OFF-sustained (A_OFF-S_) and alpha OFF-transient (A_OFF-T_) ganglion cells. A_OFF-S_ ganglion cells receive dominant inputs from type 2 (T2) and A_OFF-T_ ganglion cells are primarily driven by type 3a (T3a) OFF cone bipolar cells (Fox and Sanes, 2007; Yu et al., 2018). T2 and T3a OFF cone bipolar cells exhibit distinct temporal tuning properties (Ichinose and Hellmer, 2016) and ramify at different inner plexiform layer (IPL) depths to target A_OFF-S_ and A_OFF-T_ ganglion cells, respectively (Yu et al., 2018). Differential dynamics in these ganglion cells are thought to originate from glutamate receptors at the bipolar cell dendrites (Awatramani and Slaughter, 2000; DeVries, 2000), transmission from bipolar cells to ganglion cells, and inputs from amacrine cells (Asari and Meister, 2012; Baden et al., 2016; Franke et al., 2017). Specifically, we asked: can the responses to cone loss be localized to changes before or after the divergence of these two OFF pathways? If partial cone loss induces common changes to both A_OFF-S_ and A_OFF-T_ ganglion cells, it would indicate circuitry changes before divergence of, i.e., shared inputs to, the OFF pathways (Figure 1A). Changes could occur at one or more of the following locations: cones, horizontal cells, and/or common amacrine cells. If A_OFF-S_ and A_OFF-T_ ganglion cells exhibit differential reactions to partial cone loss, it would indicate circuitry changes after divergence of, i.e., different inputs to, the OFF pathways. Changes could occur at one or more of the following locations: bipolar cells (T2 to A_OFF-S_ and T3a to A_OFF-T_), presynaptic amacrine cells (unique ACs to T2 or T3a), and direct amacrine cells (unique ACs to A_OFF-S_ or A_OFF-T_) (Figure 1A) (Della Santina et al., 2016; Graydon et al., 2018; Yu et al., 2018).

**Figure 1.**
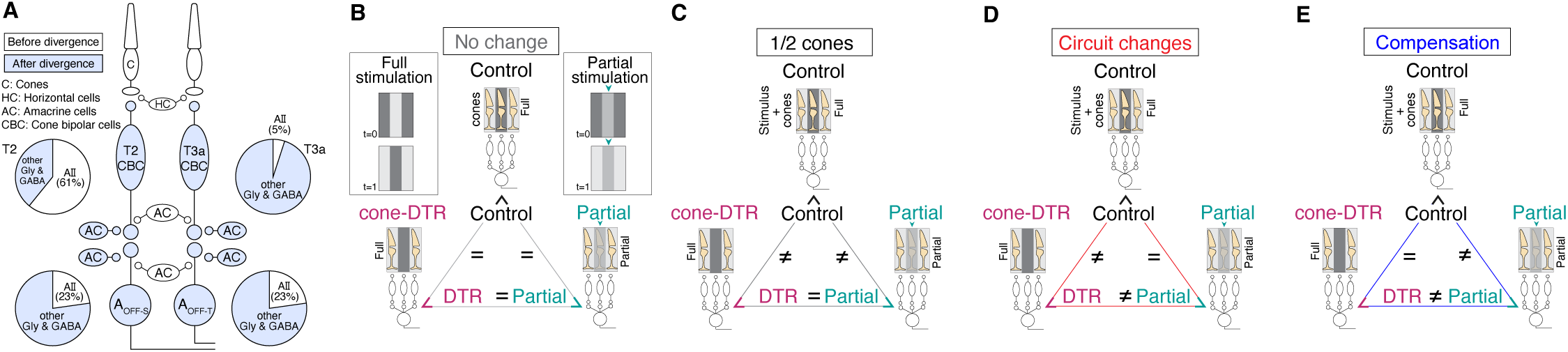
Experimental framework for distinguishing the effect of reduced cone input, circuit changes, and functional compensation following partial cone loss. (A) Illustration of known cone pathways to A_OFF-S_ and A_OFF-T_ ganglion cells. A_OFF-S_ receives approximately ≤4% inputs from type 1b and 50% inputs from type 2 (T2) cone bipolar cells (CBCs) (Della Santina et al. 2016 and Yu et al. 2018). A_OFF-T_ receives 40% inputs from T3a and 18% of inputs from T4 CBCs (Yu et al. 2018). Each CBC type receives different degrees of inhibitory inputs from AII and other amacrine cell types (top pie charts). The schematic depicts T2 and T3a CBCs as primary inputs to A_OFF-S_ and A_OFF-T_ ganglion cells, respectively. Each ganglion cell type receives glycinergic and GABAergic inhibitory inputs and equivalent contributions from AII amacrine cells (bottom pie charts). If cone ablation affects circuits before divergence of the OFF pathways, then the following sites are implicated (white): cones, horizontal cells, common amacrine cells. If cone ablation affects circuits after divergence of the OFF channels, then the following locations are implicated (blue): bipolar cells (T2 or T3a), different amacrine cells that send their inputs to sustained or transient pathways. (B-E) Comparison of three conditions and their state of cones and the retinal circuit: (1) control condition has a full complement of cones and a control circuit (control); (2) DTR condition has half the cones and a DTR circuit (cone-DTR); (3) half stimulation condition has stimulation of half the cones and a control circuit (partial). Triangles represent comparisons among three conditions that would be interpreted as (B) no change, mechanisms explained by (C) half cone stimulation, (D) circuit changes, or (E) compensation. See also Figure S2.

To examine how retinal pathways were affected by partial cone loss, we ablated a subset of cones in mature retina by injecting diphtheria toxin (DT) into mice expressing the simian DT receptor under the M-opsin promoter at postnatal day 30 (cone-DTR) (Care et al., 2019; Lee et al., 2022). The concentration of DT was adjusted to achieve 50-70% cone loss (Figure S1A-B). We examined how cone loss impacts the functional responses of OFF pathways. We targeted either A_OFF-S_ or A_OFF-T_ ganglion cells for single-cell recordings (Figure 1A). Cone-mediated inputs were elicited under three conditions: full stimulation of control retina (control), full stimulation of partial cone loss retina (cone-DTR), and partial stimulation of control retina (partial) (Figure 1B). For the partial stimulation, we modified a stimulus by blanking half the space to simulate unresponsive cones (Care et al., 2019; Lee et al., 2022). The partial stimulation was used to determine whether changes to the spatiotemporal filter induced by partial cone loss are distinct from partial stimulation, the expectation of what would occur if the retina were responding as just missing a fraction of their input with no additional circuit changes. To determine the site of changes, we used different recording configurations to isolate inputs to and outputs from A_OFF_ ganglion cells: voltage clamp to record excitatory and inhibitory input currents and current clamp to record subthreshold voltages and output spikes (Figures 2A-B, S2). We generated a linear spatiotemporal filter for each ganglion cell type in each of the three conditions under each recording configuration (Figures 2A, S2). In these three conditions, multiple aspects of the response could change. Here we focus on the linear receptive field as a key property though we acknowledge that other response aspects could change (see Methods).

**Figure 2.**
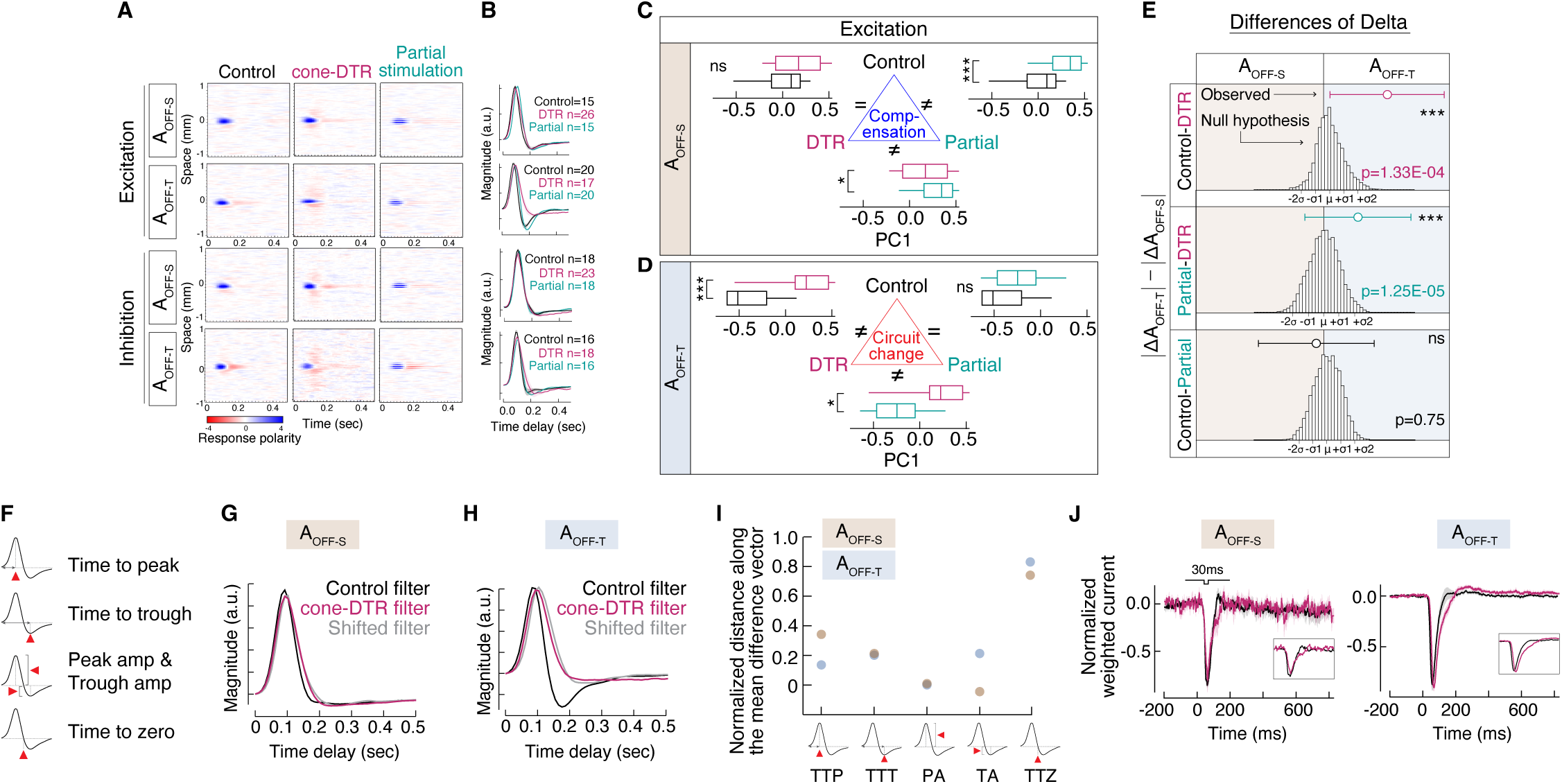
Partial cone loss causes differential temporal changes to A_OFF_ ganglion cell types. (A) Example spatio-temporal filters in response to a bar noise stimulus under three conditions for either A_OFF-S_ (rows 1, 3) or A_OFF-T_ (rows 2, 4) under voltage clamp for measuring excitation (rows 1-2) or inhibition (rows 3-4). (B) Average temporal filters of (odd rows) A_OFF-S_ and (even rows) A_OFF-T_ ganglion cells measuring excitation (rows 1-2) and inhibition (rows 3-4) for control (black), cone-DTR (magenta) and partial stimulation (green). (C-D) Box plots of first principal components (PC1) for each pairwise comparison of conditions for A_OFF-S_ and A_OFF-T_ ganglion cell excitation. Interpretation of mechanisms in the triangle center. (E) Difference of deltas results for the null hypothesis for equivalent changes in the temporal filters between A_OFF-S_ and A_OFF-T_ ganglion cells in each comparison of conditions (bootstrapped distribution of 10,000 iterations with noted standard deviations) and the actual difference of deltas (circles with error bars). Significant difference from the null hypothesis displayed on the right. (F) Features extracted from temporal filters: time to peak (TTP), time to trough (TTT), peak amplitude (PA), trough amplitude (TA), and time to zero crossing (TTZ). (G-H) Average temporal filters under control and cone-DTR conditions for (G) A_OFF-S_ and (H) A_OFF-T_ ganglion cells, which show a significant difference between control vs. cone-DTR, and a version of the control temporal filter shifted by multiple features to match the cone-DTR temporal filter. (I) Normalized distance between the temporal filter in control (ordinate = 0) and cone-DTR (ordinate = 1) when the control temporal filter is changed by each feature for A_OFF-S_ (tan) and A_OFF-T_ (blue) ganglion cells. (J) Average excitatory current normalized to the flash strength of A_OFF-S_ (left) and A_OFF-T_ (right) to a 30ms decrement in light. P-values in box plots indicate rank sum comparison between each pair of conditions after correcting for multiple comparisons with the Holm method (C-D). P-values in difference of deltas plot (E) indicate significant differences between A_OFF-S_ and A_OFF-T_ ganglion cells from a permutation test with correction for multiple comparisons. The following asterisks indicate p values: * ≤ 0.05, ** ≤ 0.01, *** ≤ 0.005. See also Figure S2 and Dataset S1.

First, we focus on the temporal component of the ganglion cell excitatory spatiotemporal filters (Figure 2B). Partial stimulation of control retina delayed the kinetics of the A_OFF-S_ ganglion cell temporal filter that is not observed in the cone-DTR retina. In contrast, following cone loss, A_OFF-T_ ganglion cells lose a biphasic shape in the excitatory temporal filter that is not observed in the control or partial stimulation conditions (Figure 2B). There are many different aspects to the temporal filter, and we do not know, *a priori,* which, if any, might change. Therefore, we performed principal components analysis of the temporal filters under all conditions (Figures S2B, C, I, O) to extract the features that vary the most, and compared the strength of the projection onto the first principal component across conditions (Figures 2C-D, S2). Given the numerous recording conditions, we present results showing significant differences between A_OFF-S_ and A_OFF-T_ ganglion cells in the main figures; the rest are in the supplementary material. To understand the mechanisms underlying changes to the temporal filter, we made pairwise comparisons of the three conditions (control vs. cone-DTR, control vs. partial stimulation, cone-DTR vs. partial stimulation) and controlled for the multiple comparisons (Figure 1B-E). For excitatory and voltage temporal filters, A_OFF-S_ ganglion cells showed differences between cone-DTR vs. partial stimulation and control vs. partial stimulation, but not between control vs. cone-DTR (Figures 2C, S2J). These results implicate compensation, i.e., a mechanism that allows responses in cone-DTR retina to match those in the full stimulation of control retina, but not match those from partial stimulation (Figure 1E). Excitatory temporal filters of A_OFF-T_ ganglion cells showed significant differences in control vs. cone-DTR and cone-DTR vs. partial, but not between control vs. partial stimulation (Figure 2D). This result implicates circuit changes, i.e., a mechanism underlying changes observed in cone-DTR that causes responses to differ from control (Figure 1D); this was also found in spike temporal filters of A_OFF-S_ ganglion cells (Figure S2P).

Next, we observed that the inhibitory temporal filter for both A_OFF-S_ and A_OFF-T_ ganglion cells in control retina had different kinetics than the cone-DTR or partial stimulation conditions (Figure 2B bottom, Figure S2D-F). In addition to the compensation and the circuit change conclusions, described above, another possible conclusion of these types of analyses would occur if there were differences in control vs. cone-DTR and control vs. partial stimulation, but not between cone-DTR vs. partial stimulation. Such a scenario would implicate half cone stimulation as the mechanism for changes observed in cone-DTR (Figure 1C). Such mechanisms were found in inhibition from both cell types and voltage temporal filters of A_OFF-T_ ganglion cells (Figure S2D-E, K).

In the pairwise comparison of the three conditions for each cell type, the identification of a difference in one cell type and not the other is insufficient to conclude differential effects between the cell types (Figure 2C-D) (Nieuwenhuis et al., 2011). Therefore, to quantify the relative differences between the two ganglion cell types, we compared the magnitude of their response changes, i.e., the difference of deltas between two cell types (see Methods). First, we created a bootstrapped distribution of the null hypothesis that A_OFF-T_ and A_OFF-S_ ganglion cells have indistinguishable changes across conditions. We then determine where the observed deltas lay relative to this distribution (Figure 2E). For two of the comparisons involving cone-DTR, i.e., (control -cone-DTR) and (partial - cone-DTR), the observed values fell in the far right tail of the null hypothesis distribution, even after controlling for the multiple comparisons, indicating that A_OFF-T_ ganglion cells undergo greater changes in the excitatory temporal filter, i.e., T3a cone bipolar cell inputs, compared to A_OFF-S_ ganglion cells (Figure 2E). Though we statistically compare all pairs of conditions, we highlight how much cone-DTR deviates from the control or partial stimulation conditions to isolate the effect of cone ablation. In contrast, the difference between control and partial conditions serves as a check of the analysis where we expect minimal difference if the observed delta is specifically driven by cone ablation. Temporal filters for subthreshold voltages showed greater changes in A_OFF-S_ ganglion cells when we compute delta between cone-DTR vs. partial stimulation but not control vs. cone-DTR (Figure S2L). For inhibitory and spike temporal filters, we found no evidence for a difference between cell types (Figure S2F, S2R). For the third comparison between full and partial stimulation of control retina, we consistently find no differences in the deltas across all conditions across recording configurations (Figures 2E, S2). Taken together, the excitatory temporal filters in the two ganglion cell types exhibited distinct mechanisms, which likely arises after the point of divergence between A_OFF-S_ and A_OFF-T_.

We next identified which features of the temporal filter changed in the excitatory currents (Figure 2F). Qualitatively, following partial cone loss, the A_OFF-S_ temporal filter was delayed (Figure 2G), while the A_OFF-T_ temporal filter lost its biphasic component (Figure 2H). Quantitatively, we measured the extent to which independently modifying each of five features, i.e., time to peak, time to trough, peak amplitude, trough amplitude, time to zero, made the control temporal filter more or less like the mean cone-DTR temporal filter. Briefly, this was achieved by linearly modifying each feature of individual control filters, and then measuring the projection of these modified filters along the vector that points from the mean control filter to the mean cone-DTR filter, where a value of 0 is equivalent to the mean control filter and a value of 1 is equivalent to the mean cone-DTR filter (see Methods, Figure S2S-U). For both cell types, time to peak, time to trough, peak amplitude, trough amplitude all contributed approximately 0.2 to 0.4 units normalized distance to the difference between control and cone-DTR temporal filters, while time to zero crossing contributed the most by shifting the temporal filter 0.8 units of normalized distance (Figure 2I). Since the analysis described above relied on the linear-nonlinear (LN) model, we quantified the accuracy of the model prediction by comparing the peak of cross-correlation between the predicted and actual responses for each recording configuration (see Methods), The same noise sequence was presented across conditions (Figure S2V) to further validate the model. These analyses did not reveal differences across conditions in either cell types. In addition, we also measured the cell’s impulse responses to decrements of light independently of the model and found both A_OFF_ cells exhibited delayed kinetics following cone loss (Figure 2J). Taken together, we conclude that following partial cone loss the changes in excitatory temporal filters of A_OFF_ ganglion cells reflect distinct mechanisms at post-receptoral sites, mostly captured by a delay in the time to zero crossing.

### Two distinct mechanisms in inhibitory circuits underlie changes in temporal filters of A_OFF-T_ ganglion cells

To reveal the circuit mechanisms underlying the temporal changes following partial cone loss, we measured excitatory and inhibitory temporal filters of A_OFF-S_ and A_OFF-T_ ganglion cells in control and cone-DTR retina in the absence and presence of picrotoxin, an antagonist of GABA_A_ and GABA_C_ receptors, or strychnine, an antagonist of glycine receptors. We analyzed temporal filters to determine the contributions of GABAergic or glycinergic inhibition.

We evaluated the effect of the GABAergic (Figure 3A, D, F, K) or glycinergic (Figure 3B, E, H, M) antagonists between each control and cone-DTR condition. To this end, we subtracted the temporal filters before and after pharmacological blockade for each cell to compute a difference filter (Figure 3C, S3A-D). Then with the difference filters, we performed principal component analysis on the control vs. cone-DTR conditions of A_OFF-S_ and A_OFF-T_ ganglion cells (Figure 3F, H, K, M, S3E-H). First, we found significant differences between the effects of glycinergic inhibition on the excitatory temporal filters of A_OFF-T_ ganglion cells under control and cone-DTR conditions (Figure 3I). Since above we found that A_OFF-T_ ganglion cells showed a change to their excitatory temporal filters (Figure 2E), these results provide a mechanism: diminished impact of presynaptic glycinergic inhibition on that circuit in the cone-DTR condition (Figure 3J).

**Figure 3.**
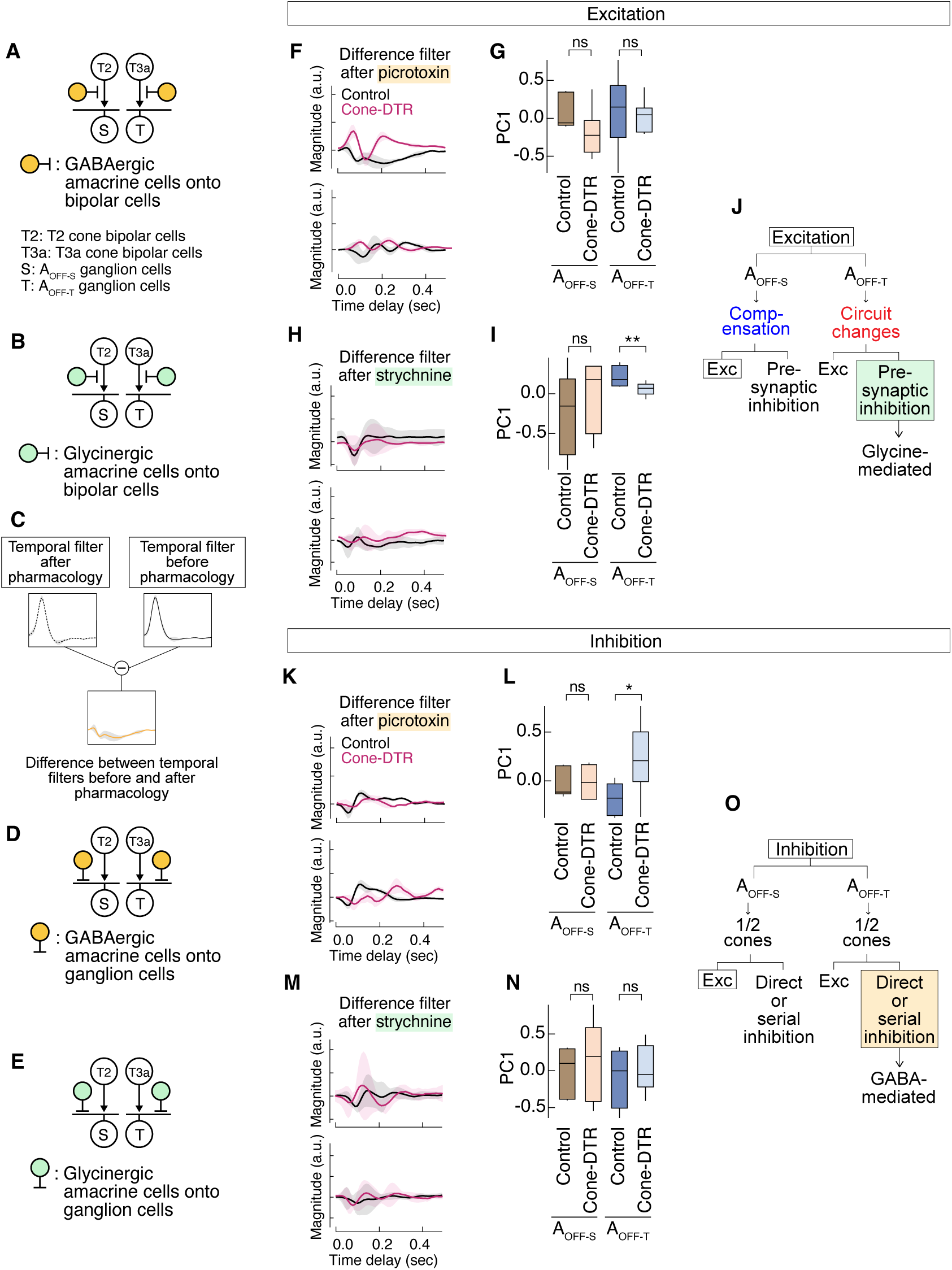
Partial cone loss causes presynaptic glycine and direct GABA inhibition to mediate changes in the cone-DTR temporal filters of A_OFF-T_ ganglion cells. (A-B, D-E) Schematic of retinal pathways leading to the A_OFF-S_ (left) and A_OFF-T_ (right) ganglion cells and the circuit motifs for direct excitation (white), presynaptic inhibition from (A) GABA (yellow) or (B) glycine (green), and direct inhibition from (D) GABA (yellow) or (E) glycine (green) that would be blocked by (A, D) picrotoxin, an antagonist of GABA_A_ and GABA_C_ receptors, or (B, E) strychnine, an antagonist of glycine receptors. (C) Illustration of the difference filter computed by subtracting the temporal filters from after and before pharmacological application for each cell. (F, H, K, M) Average difference between Ames and (F, K) picrotoxin or (H, M) strychnine. (G, I, L, N) Box plots of first principal components (PC1) across cell types and conditions for the differences between (G, L) Ames and picrotoxin or (I, N) Ames and strychnine from (G, I) excitation temporal filters and (L, N) inhibition temporal filters. (J, O) Results of the comparison of PC1 and the interpretation of mechanisms for (J) excitatory temporal filters and (O) inhibitory temporal filters. P-values in box plots indicate rank sum comparison between each pair of conditions; p-values are corrected for multiple comparisons with the Holm method (G, I, L, N). See also Figure S3 and Dataset S1.

Second, although we found no evidence for a difference between two cell types in their inhibitory temporal filters (Figure S2F), we observed significant differences between control and cone-DTR conditions under GABAergic, but not glycinergic blockade (Figure 3L, N). These results imply that increased impact of direct GABAergic input to A_OFF-T_ ganglion cells may mask underlying differences between two cell types (Figure 3O). Taken together, these data provide evidence for two modifications to circuits of the A_OFF-T_ ganglion cell’s temporal filters following partial cone loss, i.e., changes in (1) presynaptic glycinergic inhibition, consistent with changes in excitatory temporal filters (Figure 2E) and (2) direct GABAergic inhibition, which could arise from loss of cone inputs (Figure S2C-F).

### Partial cone loss causes narrowing of center and widening of receptive field surround in A_OFF_ ganglion cells

Having observed changes to the temporal dimension of the ganglion cell responses, we next determined how partial cone loss affected spatial processing of these OFF pathways. We applied the same framework as described for the kinetics to determine the contributions of half cone stimulation, circuit changes, and compensation to the spatial filters of each cell type (Figure 1B-E, S4A-B); to compare spatial filter differences between two cell types; and to determine if these mechanisms arise prior to or after divergence between the OFF pathways.

In excitatory spatial receptive fields, A_OFF-T_ ganglion cells exhibit a change in cone-DTR distinct from the control and partial stimulation conditions (Figure 4B-C). In the voltage spatial receptive fields, both A_OFF-S_ and A_OFF-T_ ganglion cells show significant changes in the cone-DTR condition (Figure 4G-I). To evaluate these changes across conditions, principal component analysis was performed on the entire spatial filter for excitation, inhibition, spikes, and subthreshold voltages (Figures S4C). The projection onto the first principal component of the spatial filters were significantly different between cone-DTR vs. control and cone-DTR vs. partial conditions for A_OFF-T_ excitation (Figure 4B) and voltages for both A_OFF-S_ and A_OFF-T_ voltages (Figure 4G-H), which we attribute to circuit changes (Figure 1D). In contrast, the projection onto the first component of the A_OFF-T_ inhibitory spatial filters (Figure S4D) and A_OFF-S_ spike spatial (Figure S4F) were significantly different between cone-DTR vs. partial and control vs. partial, which we attribute to compensation (Figure 1E). Greater spatial changes to excitatory spatial filters of A_OFF-T_ (Figure 4C) and voltage spatial filters of A_OFF-S_ ganglion cells (Figure 4I) implies distinct circuit changes in post-receptoral circuits, i.e., cone bipolar cells, unique amacrine cell inputs, and/or intrinsic properties of ganglion cells (Figure 1D).

**Figure 4.**
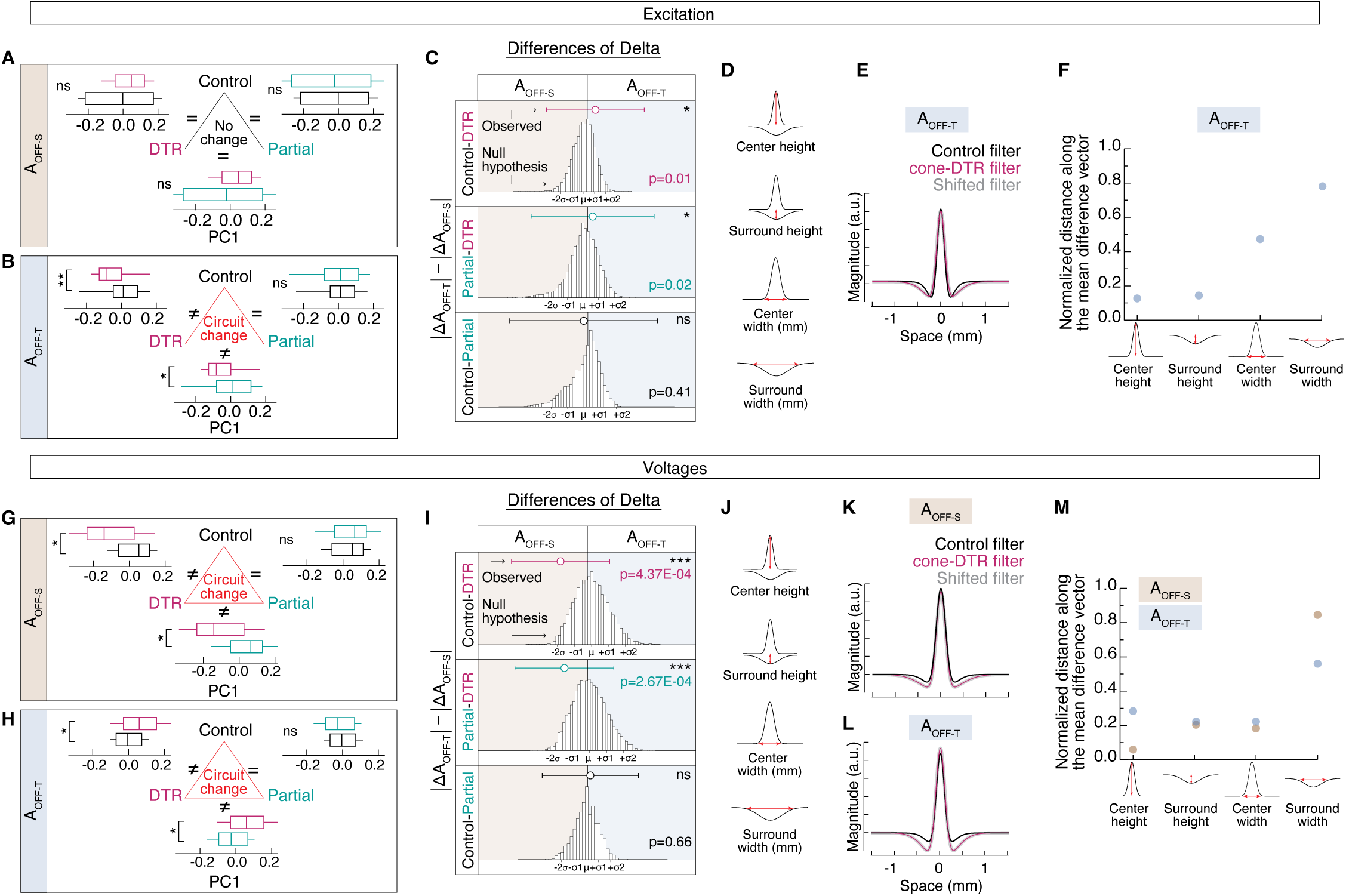
Partial cone loss causes differential changes in spatial filters of A_OFF_ ganglion cell types. (A-B, G-H) Box plots of first principal components (PC1) for (A, G) A_OFF-S_ and (B, H) A_OFF-T_ ganglion cells between conditions from (A-B) excitation and (G-H) voltages, which showed significant differences across conditions. Interpretation of mechanisms in the triangle centers. (C, I) Difference of deltas results for the null hypothesis for equivalent changes in the spatial filters between A_OFF-S_ and A_OFF-T_ ganglion cells in each comparison of conditions (bootstrapped distribution of 10,000 iterations with noted standard deviations) and the actual difference of deltas (circles with error bars) for (C) excitation and (I) voltages. Significant difference from the null hypothesis displayed on the right. (D, J) Features extracted from spatial filters: center height, surround height, center width, and surround width for (D) excitation and (J) voltages. (E, K-L) Average spatial filters under control and cone-DTR conditions for (E) A_OFF-T_ ganglion cell excitation and (K) A_OFF-S_ and (L) A_OFF-T_ ganglion cell voltages, which show a significant difference between control vs. cone-DTR, and a version of the control spatial filter shifted by multiple parameters to match the cone-DTR spatial filter. (F, M) Normalized distance between the spatial filter in control (ordinate = 0) and cone-DTR (ordinate = 1) when the control spatial filter is changed by each parameter for (F) A_OFF-T_ ganglion cell excitation and (M) A_OFF-S_ and A_OFF-T_ ganglion cell voltages. P-values in box plots indicate rank sum comparison between each pair of conditions; p-values are corrected for multiple comparisons with the Holm method (A-B, G-H). P-values in the difference of deltas plots indicate significant differences between A_OFF-S_ and A_OFF-T_ ganglion cells from a permutation test with correction for multiple comparisons (C, I). See also Figure S4 and Dataset S1.

Similar to the way we analyzed the temporal filters, we next determined which features of the spatial filter changed in the excitation of A_OFF-T_ ganglion cells and voltages of both A_OFF_ ganglion cells (Figure 4D-E, J-L) by assessing how modifying each feature of the control spatial filter better matches the cone-DTR filter. Qualitatively, for the excitation of A_OFF-T_ ganglion cells following partial cone loss, the center of the spatial filter narrowed (Figure 4E). Quantitatively, while center and surround height contribute approximately 0.1 units to the difference between control (normalized value of 0) and cone-DTR (normalized value of 1) spatial filters, center and surround widths contribute a shift of 0.4 and 0.8 units toward cone-DTR spatial filters, respectively (Figure 4F). Qualitatively, for the voltages of both A_OFF_ ganglion cells following partial cone loss, the surround of the spatial filter widened (Figure 4K-L). Quantitatively, center and surround height and center width each contribute approximately 0.1-0.3 units to the difference between control (normalized value of 0) and cone-DTR (normalized value of 1) spatial filters (Figure 4M). While the surround width of A_OFF-T_ contributes a shift of 0.55 units toward cone-DTR spatial filters, that of A_OFF-S_ contributes 0.85 units, consistent with greater changes in A_OFF-S_ compared to A_OFF-T_ ganglion cells (Figure 4I). Taken together, following partial cone loss, the changes in spatial filters from excitation of A_OFF-T_ and voltage of both A_OFF_ ganglion cells reflect circuit mechanisms mostly captured by a narrowing of the center and widening of the surround. These changes could arise from loss of cone inputs to bipolar cells and/or changes in presynaptic inhibition to bipolar cell inputs, and changes in amacrine cell inputs to both A_OFF_ ganglion cells independently.

### Two distinct mechanisms underlie changes in spatial filters of A_OFF-T_ ganglion cells

Next, we determined the mechanisms underlying the spatial changes following partial cone loss. We measured excitatory and inhibitory spatial filters of A_OFF-S_ and A_OFF-T_ ganglion cells in control and cone-DTR retina in the absence and presence of an antagonist of GABA_A_ and GABA_C_ receptors (Figure 5A, D, F, K), or an antagonist of glycine receptors (Figure 5B, E, H, M). With the difference filters of the spatial filters before and after pharmacological blockade for each cell (Figure 5C, S3I-L), we performed principal component analysis on the control and cone-DTR conditions of A_OFF-S_ and A_OFF-T_ ganglion cells (Figure 5F, H, K, M, S3M-P). First, for the A_OFF-S_ ganglion cells, we found no differences in the first principal components of the spatial filters between control vs. cone-DTR conditions under both GABAergic and glycinergic blockade (Figure 5G, I); therefore, we attributed the mechanism underlying the changes in cone-DTR in the excitatory spatial filters of A_OFF-S_ ganglion cells to excitatory circuits (Figure 5J). Second, for the A_OFF-T_ ganglion cells, we found a significant difference in the first principal components of the spatial filters between control vs. cone-DTR conditions under glycinergic blockade (Figure 5I). Since we found that A_OFF-T_ ganglion cells showed a difference in the changes to their excitatory spatial filters (Figure 4C), these results suggest that the observed difference may result from diminished presynaptic glycinergic inhibition onto bipolar cells in the cone-DTR condition (Figure 5J). Finally, similar to inhibitory temporal filters, we found no evidence for a difference between two cell types in their inhibitory spatial filters (Figure S4E). However, we observed significant differences between control and cone-DTR conditions under GABAergic, but not glycinergic blockade (Figure 5L, N). These results imply that increased impact of direct GABAergic input to A_OFF-T_ ganglion cells may mask underlying differences between two cell types (Figures 5O, S4E).

**Figure 5.**
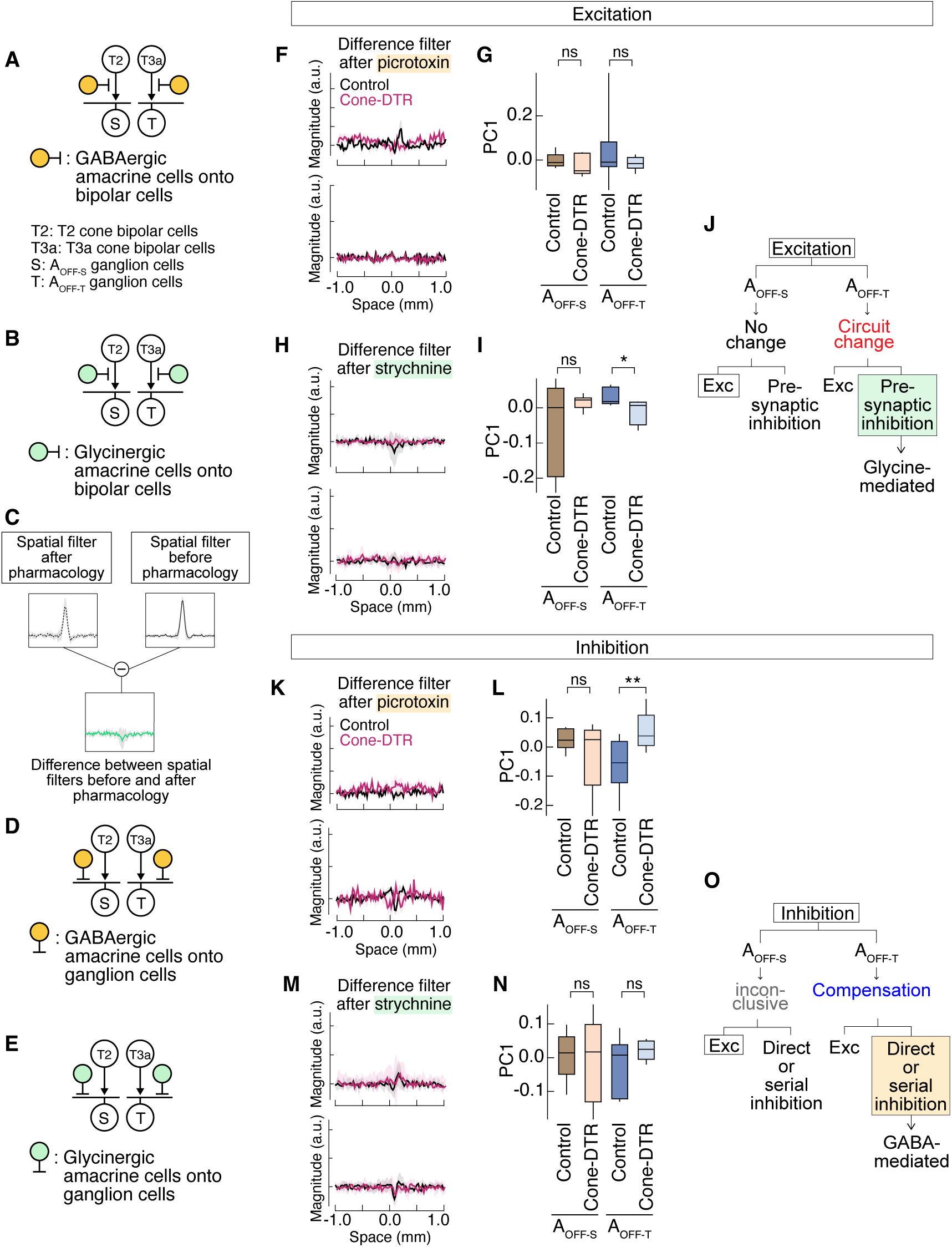
Partial cone loss causes excitation and direct GABAergic and glycinergic inhibition to mediate changes in the cone-DTR spatial filters of A_OFF-T_ ganglion cells. (A-B, D-E) Schematic of retinal pathways leading to the (left) A_OFF-S_ and (right) A_OFF-T_ ganglion cells and the circuit motifs for direct excitation (white), presynaptic inhibition from (A) GABA (yellow) or (B) glycine (green), and direct inhibition from (D) GABA (yellow) or (E) glycine (green) that would be blocked by (A, D) picrotoxin, an antagonist of GABA_A_ and GABA_C_ receptors, or (B, E) strychnine, an antagonist of glycine receptors. (C) Illustration of the difference filter computed from subtracting the spatial filters from after and before pharmacological application for each cell. (F, H, K, M) Average difference between Ames and (F, K) picrotoxin or (H, M) strychnine. Variance captured by each principal component noted on the corresponding axis. (G, I, L, N) Box plots of first principal components (PC1) across cell types and conditions for the differences between (G, L) picrotoxin and Ames or (I, N) strychnine and Ames from (G, I) excitation spatial filters and (L, N) inhibition spatial filters. (J, O) Results of the comparison of PC1 and the interpretation of mechanisms for (J) excitatory spatial filters and (O) inhibitory spatial filters. P-values in box plots indicate rank sum comparison between each pair of conditions; p-values are corrected for multiple comparisons with the Holm method (G, I, L, N). See also Figure S3 and Dataset S1.

While modifications in the inhibitory spatial filters of A_OFF-S_ ganglion cells have two possible mechanistic interpretations (Figure 5O), the effects of pharmacology on the spatial filters provide evidence for three modifications to circuits of the A_OFF_ ganglion cells following partial cone loss, i.e., changes in (1) excitation to A_OFF-S_, and (2) presynaptic glycinergic and (3) direct GABAergic inhibition to A_OFF-T_ ganglion cells.

### Partial cone loss affects nonlinearities at multiple levels leading to partial recovery of A_OFF_ ganglion cell outputs

The linear spatiotemporal filters described above map onto the cell’s actual response with a nonlinearity, which reflects average changes in response magnitude to a given change in stimulus contrast. For both cell types, cone-DTR shows shallower input nonlinearities, compared to control, that become steeper in output voltages and spikes (Figure 6A). To quantify this input-to-output transformation, we performed principal component analysis on the entire nonlinearity, where more negative PC1 corresponds to a shallower nonlinearity (Figure S5A). Indeed, we found that cone-DTR exhibited decreased excitatory input currents to A_OFF-S_ and A_OFF-T_ (Figure 6B-C), and inhibitory input currents to A_OFF-T_ (Figure 6F) ganglion cells compared to both full and partial stimulation of control conditions. Therefore, we attribute the mechanism to circuit changes, which arise from bipolar and amacrine cells. For the subthreshold voltage nonlinearities, the A_OFF-S_ ganglion cells had no significant differences in the cone-DTR vs. control, i.e., no change (Figure 6H). In contrast, A_OFF-T_ ganglion cells showed significantly shallower voltage nonlinearities in cone-DTR compared to control (Figure 6I). This demonstrates that the nonlinearity does not fully recover to control values, consistent with greater changes observed in A_OFF-T_ ganglion cells in cone-DTR retina (Figure 6J). Interestingly, although the mechanism of voltages of A_OFF-T_ remains inconclusive (Figure 6I), spike nonlinearities in the cone-DTR condition were significantly different from both control and partial stimulation, i.e., with a mechanistic interpretation of circuit changes (Figure 6L). This results in greater changes to A_OFF-T_ compared to A_OFF-S_ ganglion cells (Figure 6M). Unlike the reduced inputs, the PC1 of spike nonlinearities in A_OFF-T_ ganglion cells of cone-DTR shifted toward positive PC1 values, i.e., steeper nonlinearity, demonstrating recovery of the voltage-to-spike transformation (Figure 6L).

**Figure 6.**
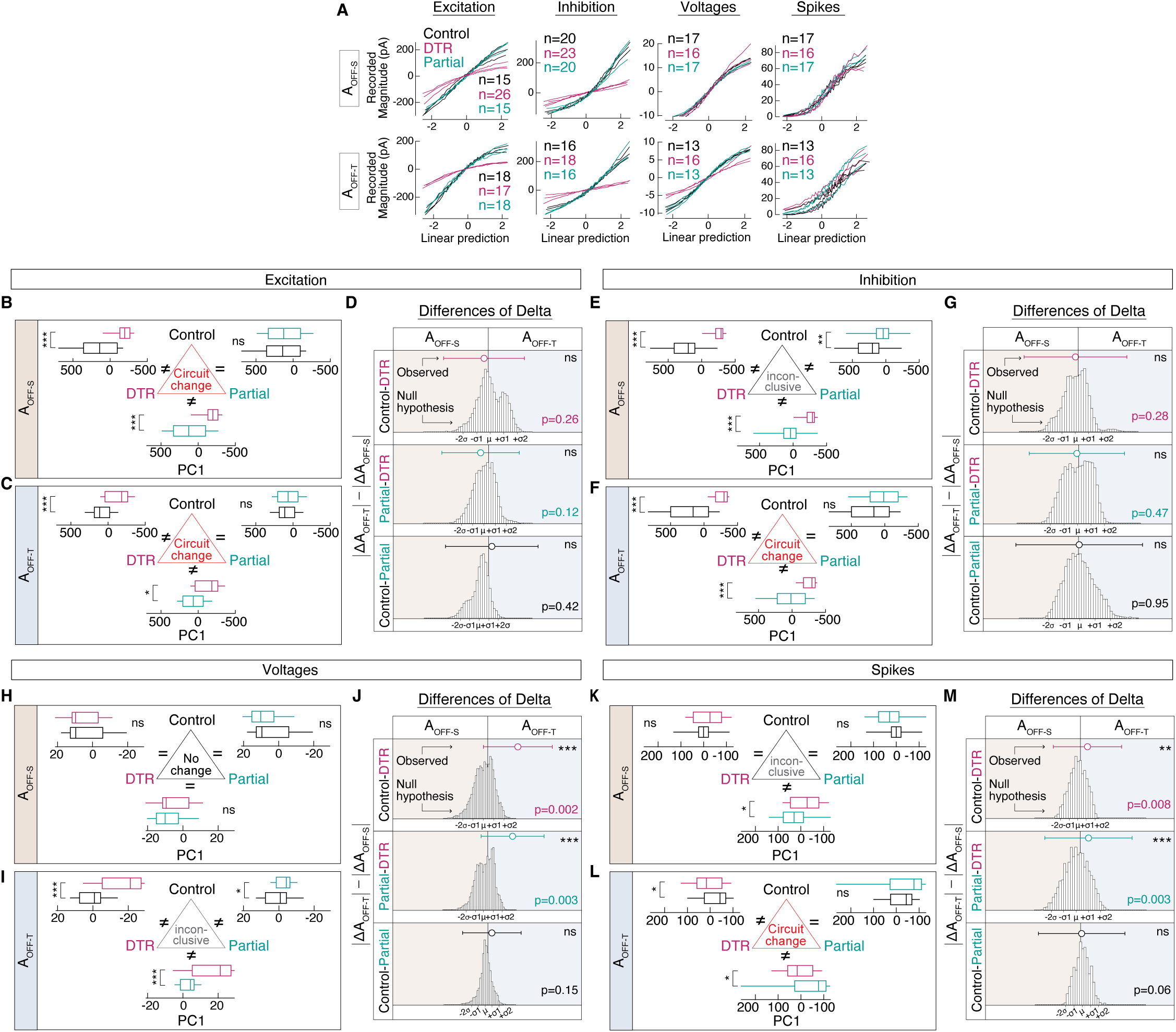
Partial cone loss affects nonlinearities at multiple levels leading to partial recovery of A_OFF_ ganglion cell outputs. (A) Three exemplar nonlinearities from control (black), cone-DTR retina (magenta), and partial stimulation (green) for excitation, inhibition, subthreshold voltages, and spikes from individual (top) A_OFF-S_ and (bottom) A_OFF-T_ ganglion cells. (B-C, E-F, H-I, K-L). Box plots of first principal components (PC1) for A_OFF-S_ and A_OFF-T_ ganglion cells between conditions from (B-C) excitation, (E-F) inhibition, (H-I) voltages, (K-L) spikes. Interpretation of mechanisms in the triangle centers. (D, G, J, M) Difference of deltas results for the null hypothesis for equivalent changes in the nonlinearity between A_OFF-S_ and A_OFF-T_ ganglion cells in each comparison of conditions (bootstrapped distribution of 10,000 iterations with noted standard deviations) and the actual difference of deltas (circles with error bars) for (D) excitation, (G) inhibition, (J) voltages, and (M) spikes. Significant difference from the null hypothesis displayed on the right. P-values in box plots indicate rank sum comparison between each pair of conditions; p-values are corrected for multiple comparisons with the Holm method (B-C, E-F, H-I, K-L). P-values in the difference of deltas plots from a permutation test with correction for multiple comparisons (D, G, J, M). See also Figures 7, S5 and Dataset S1.

Having found these differences between input and output nonlinearities, we next evaluated whether cone loss differentially affects nonlinearities of inputs vs. outputs across conditions. In both cell types, cone loss consistently resulted in greater changes in input currents than in output voltages or spikes (Figure 7B, D, F, H). To verify the quantification method, the comparisons between control vs. partial showed no significant differences between input currents and output voltages and spikes as expected.

**Figure 7.**
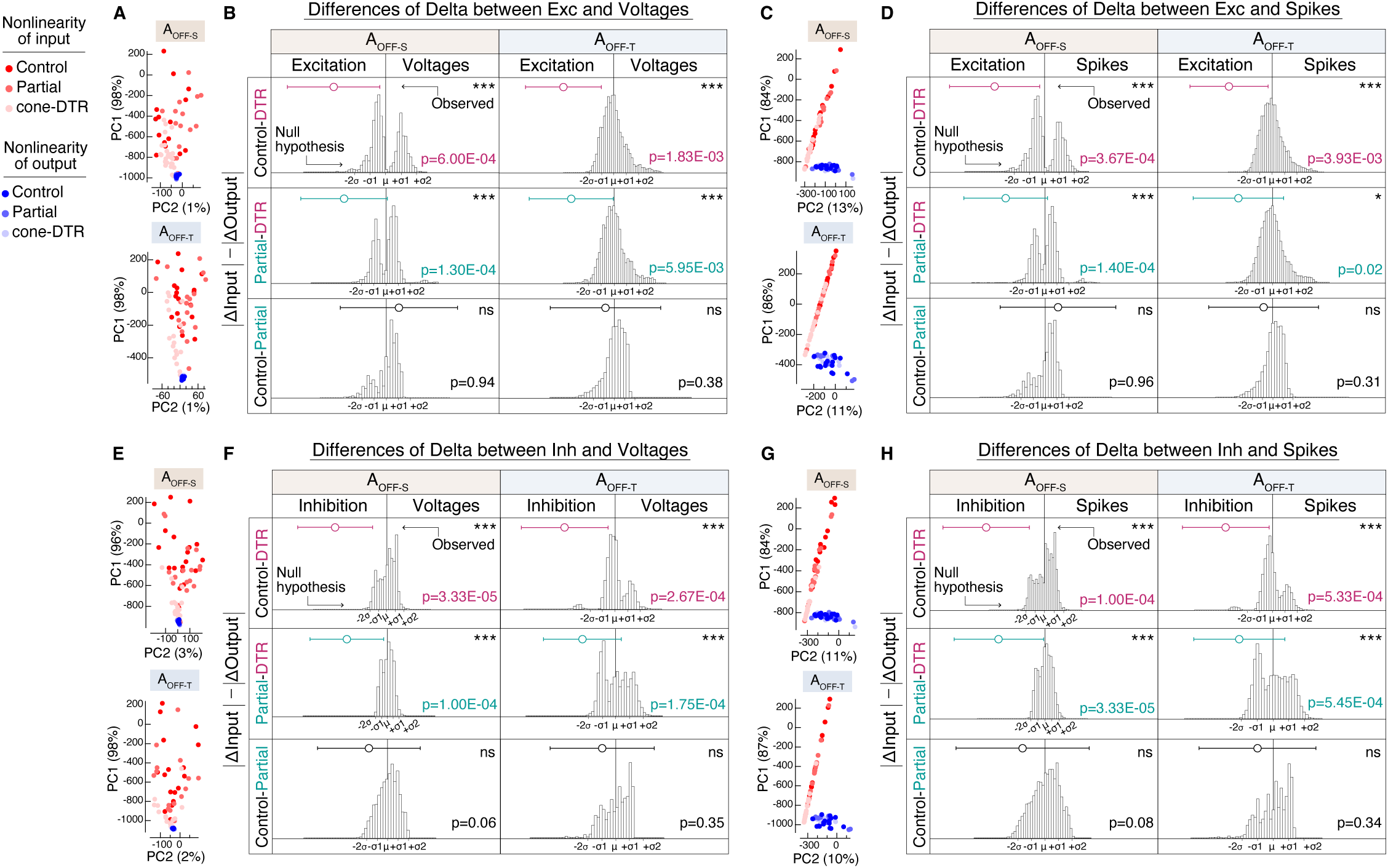
Partial cone loss has a differential impact on nonlinearities of input currents vs. output voltages and spikes. (A, C, E, G) Principal component analysis was performed on the nonlinearities for (A) excitation and subthreshold voltages, (D) excitation and spikes, (E) inhibition and subthreshold voltages, (G) inhibition and spikes. (B, D, F, H) Using the projection onto the first principal component of nonlinearities across conditions, we analyzed the difference of deltas to determine differences between inputs vs. outputs within each cell type. P-values in the difference of deltas plots from a permutation test with correction for multiple comparisons (B, D, F, H). See also Dataset S1.

Given that the spike nonlinearities of cone-DTR were different from control (Figure 6L-M), we asked whether cone loss induced changes in intrinsic excitability of A_OFF-T_ ganglion cells. We tested this by measuring spikes in response to injected white noise currents into A_OFF_ ganglion cells under perforated patch-clamp configuration (Chichilnisky and Kalmar, 2002; Care et al., 2019; Lee et al., 2022). We found no evidence to support this possibility for both A_OFF-S_ and A_OFF-T_ ganglion cells (Figure S5B-G). However, changes in intrinsic properties, such as resting membrane potential and spike threshold, could be missed as the membrane potentials were artificially held at -60mV. Therefore, we injected a slow ramp of current to induce spikes to capture intrinsic differences between these two cell types. This revealed that cone loss depolarized the resting membrane potential of A_OFF-T_ ganglion cells with a maintained spike threshold (Figure S5H-I), which could account for a steeper voltage-to-spike transformation (Figure 6A). Although the effect of cone-DTR resulted in inconclusive mechanisms in inhibitory currents from A_OFF-S_ ganglion cells (Figure 6E), shallower nonlinearities of input currents in both cell types suggest common effects of cone loss on two OFF pathways. This was further supported by the greater effect of cone-DTR onto input currents than outputs. Furthermore, nonlinearity differences between subthreshold voltages and spikes arise from intrinsic differences in A_OFF-T_ ganglion cells, suggesting that in addition to cone loss affecting cone bipolar and amacrine cells, cone loss evoked changes at the level of the ganglion cells.

### Three distinct mechanisms underlie changes in nonlinearities of both A_OFF-S_ and A_OFF-T_ ganglion cells upon partial cone loss

Having found circuit changes to excitatory and inhibitory nonlinearities following partial cone loss (Figures 6 and 7), we evaluated the differences between the input current nonlinearities before and after GABAergic or glycinergic antagonists (Figure 8G, I, L, N, S3Q-X) to identify underlying mechanisms. The first principal components of the excitatory nonlinearity differences were not significantly different between control vs. cone-DTR conditions for both GABAergic and glycinergic blockade, suggesting changes in excitatory circuits to A_OFF-S_ and A_OFF-T_ ganglion cells of cone-DTR retina (Figure 8F-J). On the other hand, we found the first principal components of inhibitory nonlinearity differences from both ganglion cell types were significantly different between control vs. cone-DTR conditions under GABAergic blockade (Figure 8L), suggesting an increased impact of direct GABAergic inhibition onto A_OFF-S_ and A_OFF-T_ ganglion cells (Figure 8O). In addition, the first principal components of the inhibitory nonlinearity differences in A_OFF-T_ ganglion cells from cone-DTR had an increased effect of glycinergic blockade from that of control retina (Figure 8N). Therefore, we attributed the mechanism underlying changes to the nonlinearity of A_OFF-T_ ganglion cells to both direct GABAergic and glycinergic inhibition (Figure 8O).

**Figure 8.**
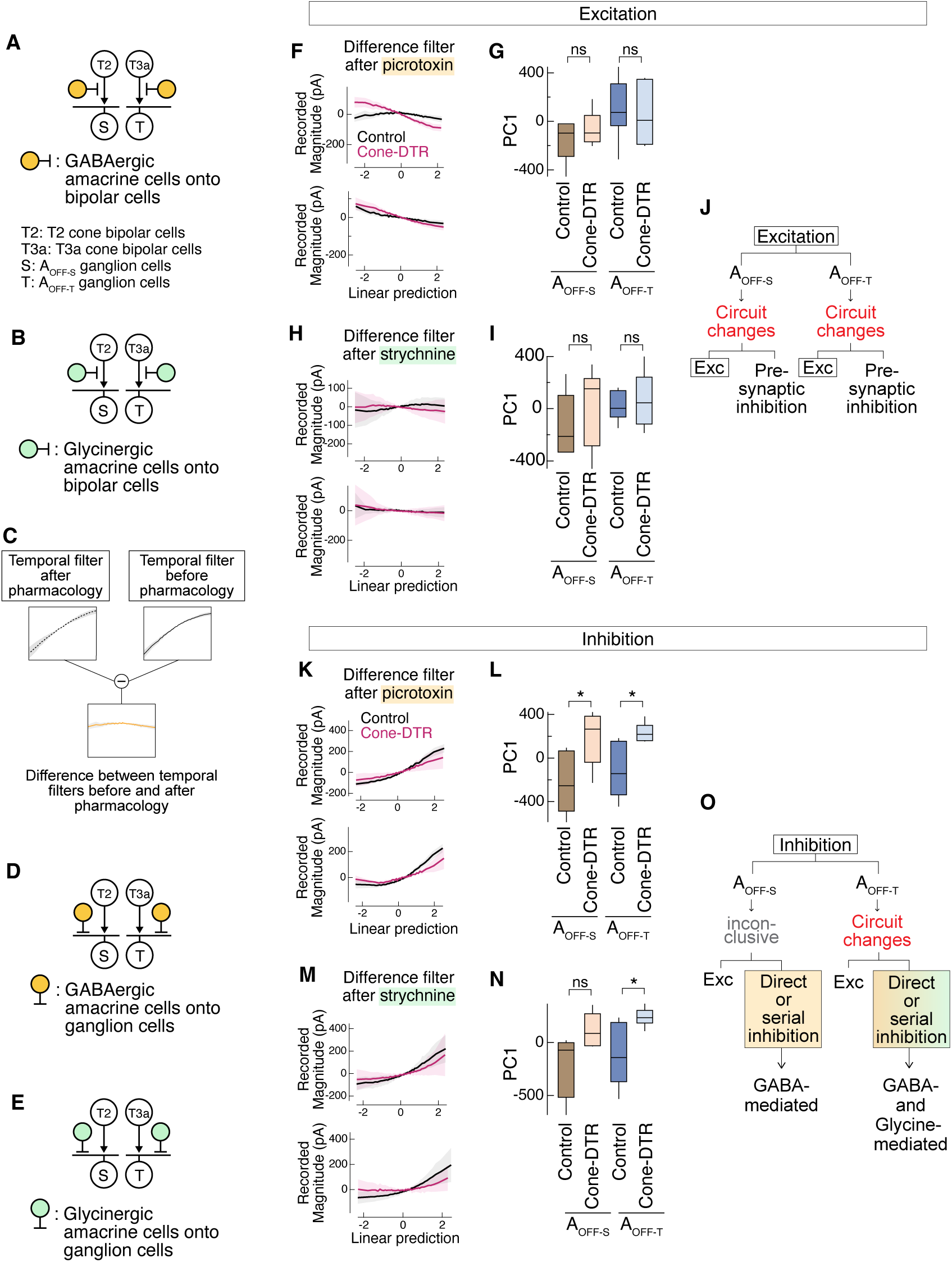
Partial cone loss causes excitation, direct GABA and glycine inhibition to mediate changes in the cone-DTR nonlinearities of A_OFF_ ganglion cells. (A-B, D-E) Schematic of retinal pathways leading to the (left) A_OFF-S_ and (right) A_OFF-T_ ganglion cells and the circuit motifs for direct excitation (white), presynaptic inhibition from (A) GABA (yellow) or (B) glycine (green), and direct inhibition from (D) GABA (yellow) or (E) glycine (green) that would be blocked by (A, D) picrotoxin, an antagonist of GABA_A_ and GABA_C_ receptors, or (B, E) strychnine, an antagonist of glycine receptors. (C) Illustration of the difference nonlinearity computed from subtracting the nonlinearities from after and before pharmacological application for each cell. (F, H, K, M) Average difference between Ames and (F, K) picrotoxin or (H, M) strychnine. (G, I, L, N) Box plots of first principal components (PC1) across cell types and conditions for the differences between (G, L) picrotoxin and Ames or (I, N) strychnine and Ames from (G, I) excitation nonlinearities and (L, N) inhibition nonlinearities. (J, O) Results of the comparison of PC1 and the interpretation of mechanisms for (J) excitatory and (O) inhibitory nonlinearities. P-values in box plots indicate rank sum comparison between each pair of conditions; p-values are corrected for multiple comparisons with the Holm method (G, I, L, N). See also Figure S3 and Dataset S1.

Having found functional evidence for glutamatergic, GABAergic, and glycinergic modifications following partial cone loss, we examined synaptic puncta in the bipolar and ganglion cells of these two retinal pathways. We found a subset of the anatomical parameters correlated with functional findings (Figure 9). Specifically, we found increased size of excitatory synapses at T2 and T3a bipolar cells (Figure 9F-G) and increased density of GABA_A_ receptors at T3a bipolar cells (Figure 9E). Also we found increased size of GABA_A_ receptors at both A_OFF_ ganglion cell types (Figure 9K-L).

**Figure 9.**
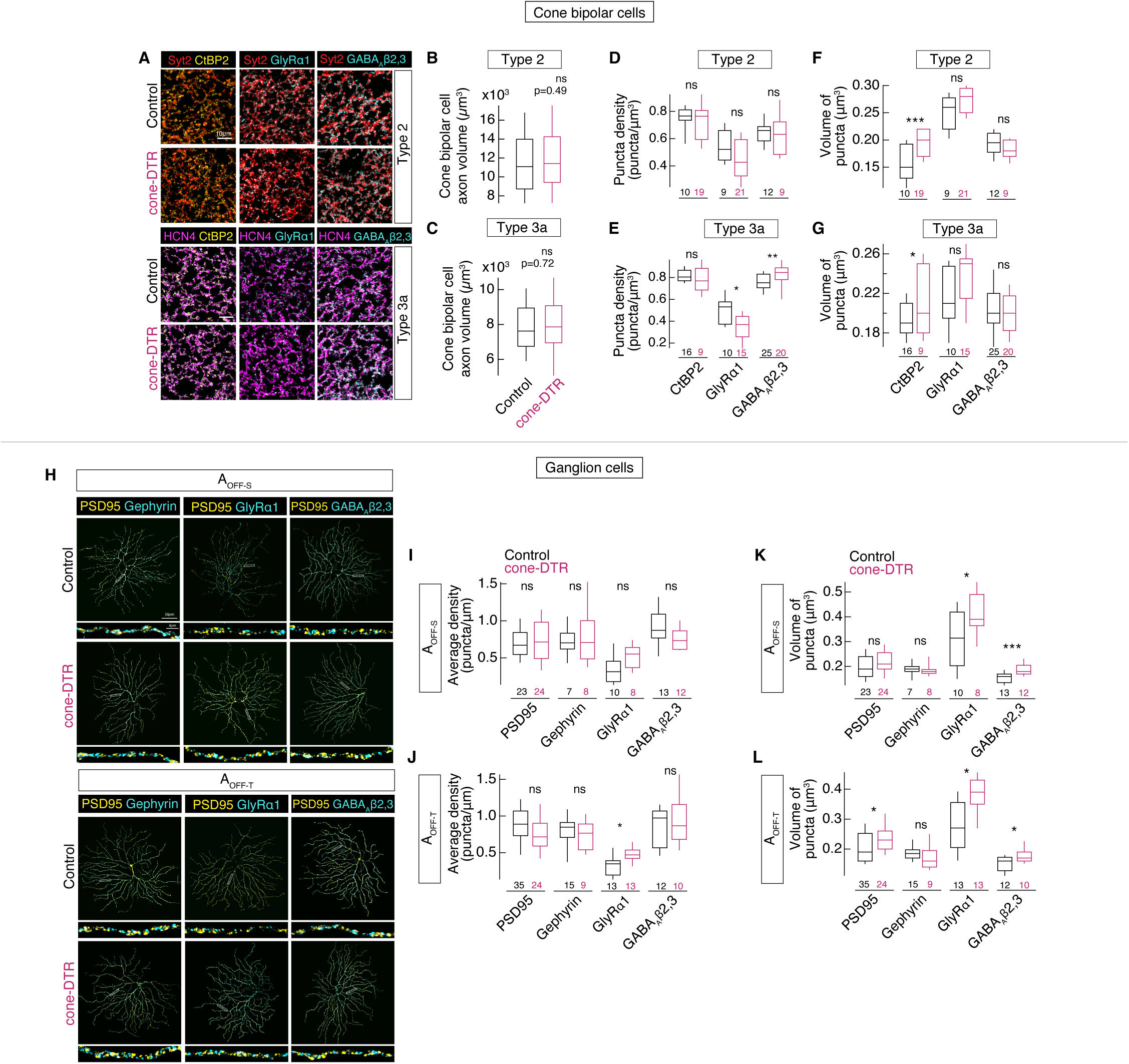
Excitatory and inhibitory synapses change in OFF pathways following partial cone loss. (A) *En face* confocal images of T2 (rows 1-2) and T3a (rows 3-4) OFF CBC axons as labeled by synaptotagmin 2 (Syt2) and HCN4 (magenta) with CtBP2 (yellow) for excitatory output synapses, GlyRɑ1 for glycine receptor alpha-1, and GABA_A_β2,3 (cyan) for GABA_A_ receptor subunit beta 2-3 in control (rows 1, 3) and cone-DTR (rows 2, 4) retina. (B-C) The volume of cone bipolar cell axon terminals in cone-DTR retina remained constant. (D-G) While the puncta density of ribbon synapses was maintained in the axons of both cone bipolar cell populations (D-E), we found an increased volume of CtBP2 in both bipolar cell types (F-G). In contrast to the enlarged volume of excitatory synapses, that of inhibitory synapses remained unchanged. Interestingly, we found differential changes in the density of GlyRɑ1 and GABA_A_Rβ2,3 in T3a cone bipolar cell (D-E). These results demonstrate common adjustments of excitatory synaptic ribbons in response to partial cone loss whereas inhibitory synapses potentially undergo differential changes between T2 and T3a cone bipolar cells. (H) *En face* confocal images of control A_OFF-S_ (rows 1-2) and A_OFF-T_ (rows 3-4) ganglion cells labeled with PSD95, GlyRɑ1, gephyrin, and GABA_A_β2,3 and in cone-DTR retina. Insets show a stretch of dendrites with puncta for rectangles in main images. (I-J) Average puncta volume of PSD95, gephyrin, GlyRɑ1, and GABA_A_β2,3 within the dendrites of (I) A_OFF-S_ and (J) A_OFF-T_ ganglion cells. While the volume and density of PSD95 remained constant in A_OFF-S_ ganglion cells, A_OFF-T_ ganglion cells showed increased PSD95 puncta size (I-J). We also found increased volume of GlyRɑ1 and GABA_A_Rβ2,3 clusters in both ganglion cell types (I-J). (K-L) Average linear density of PSD95, gephyrin, GlyRɑ1, and GABA_A_β2,3 within the dendrites of (K) A_OFF-S_ and (L) A_OFF-T_ ganglion cells. Despite a common increase in the volume of inhibitory receptors, A_OFF_ ganglion cells exhibit differential changes to synaptic density (K-L). Density and volume of gephyrin were maintained in both cell types in cone-DTR retina. Box plots show median with IQR and whiskers from 10% to 90% of the data. P-values indicate rank sum comparison between cell types. See also Dataset S1.

Taken together, these data provide evidence for common modifications of excitatory circuits following partial cone loss in A_OFF-S_ and A_OFF-T_ ganglion cells. Additionally, inhibitory circuits undergo two distinct modifications: (1) direct GABAergic inhibition onto A_OFF-S_ ganglion cells, and (2) direct GABAergic and glycinergic inhibition onto A_OFF-T_ ganglion cells.

## DISCUSSION

Results demonstrate that both A_OFF-S_ and A_OFF-T_ ganglion cells undergo different temporal and spatial modifications in response to partial cone loss that cannot be recapitulated by partial stimulation of control retina, providing evidence for changes unique to each pathway. We categorized these changes in the following mechanisms: A_OFF-S_ ganglion cells undergo compensatory changes and A_OFF-T_ ganglion cells undergo circuit changes which lead to differences in their linear temporal filters (Figure 2). Pharmacological manipulations revealed changes to excitatory and inhibitory circuits of A_OFF-T_ ganglion cells, i.e., presynaptic glycinergic inhibition and direct GABAergic inhibition (Figure 3). For A_OFF-S_ ganglion cells, the absence of effects for blockade of glycine and GABA receptor antagonists suggests potential compensation in the excitatory temporal filters, which could be attributed to changes in direct excitatory input; however, we leave open the possibility that serial inhibition cancels the effects of each antagonist applied independently (Figure 3O). We also found that changes in excitatory and inhibitory spatial filters arise from glycinergic presynaptic inhibition and direct GABAergic inhibition onto A_OFF-T_ ganglion cells (Figures 4-5). Lastly, unlike input nonlinearities, we found recovery of the voltage-to-spike transformation in A_OFF-T_ ganglion cells, which involves changes in intrinsic properties (Figures 6, S5H). Pharmacological manipulations revealed mechanistic changes to A_OFF-S_ and A_OFF-T_ ganglion cells, i.e., excitation and direct GABAergic and glycinergic inhibition (Figure 8), suggesting that cone loss affects nonlinearities at multiple levels that ultimately lead to partial recovery of A_OFF-T_ ganglion cell outputs. While A_OFF-S_ ganglion cells receive direct inhibitory inputs from AII amacrine cells (Manookin et al., 2008; van Wyk et al., 2009), we did not detect changes in glycinergic inhibition in A_OFF-S_ cells across conditions. In contrast, we found changes in both glycinergic and GABAergic inhibition onto A_OFF-T_ ganglion cells indicating modifications of multiple inhibitory pathways, potentially including AII amacrine cells. Each of these results support the conclusions that neither partial light stimulation alone nor one common mechanism is sufficient to elicit the changes following partial cone loss. Our study finds multiple sites of modifications in the retinal circuit after divergence of the two OFF pathways, indicating functional differences in response to common input loss.

Previously, we showed that partial cone loss triggers changes in inhibitory circuits in the ON pathway, specifically in pathways leading to the A_ON-S_ ganglion cells. A_ON-S_ ganglion cells exhibited expansion of receptive field surrounds, greater temporal integration, recovery of inhibitory gain and enhancement of inhibitory synapses in the dendrites. We determined that inhibition compensates for fewer cones to preserve perception of static natural images (Lee et al., 2022). Both ON and OFF pathways exhibited some common spatial changes following partial cone loss, i.e., surround expansion. Yet, we found a difference in adjusting kinetics across retinal pathways after cone loss (Figure 2) consistent with previous studies in control retina (Chander and Chichilnisky, 2001; Kim and Rieke, 2001). Specifically, cone-DTR retina of A_OFF-T_ ganglion cells exhibited greater degree of changes in excitatory temporal filters compared to that of A_OFF-S_ ganglion cells. Expansion of the surround and loss of the biphasic component could be attributed to potential changes in inhibition, for example, changes in inhibitory inputs onto A_OFF-T_ ganglion cells (Figures 3, 5, 8 and 9). At the same time, we found that changes in excitatory circuits are a common consequence of global disruption of cones. Future experiments assessing the potential upregulation of developmentally-expressed receptor subunits following cone loss will determine the contributions of GABAergic and glycinergic amacrine cells in shaping visual outputs (Sinha et al., 2021). A_OFF-T_ ganglion cells tend to be more vulnerable among the mouse ganglion cell types in other manipulations of the retinal circuit, e.g., models of glaucoma and optic nerve crush (Ou et al., 2016; Tran et al., 2019a). Perhaps differences in manipulation cause differential responses across ganglion cell types and/or perhaps the plasticity of inhibitory circuits leading to the A_OFF-T_ ganglion cells render them robust in some conditions or up to certain thresholds of damage and then render them particularly vulnerable beyond those thresholds. Future work comparing how retinal circuits react across manipulations could distinguish among these possibilities. Finally, considering the transcriptional similarity between A_OFF-T_ and A_OFF-S_ ganglion cell types (Tran et al., 2019a), differences in vulnerability of these two cell types across manipulations could be attributed to properties not discernible by their transcriptomic profiles and/or by their extrinsic properties, i.e., circuit inputs.

Spatial and temporal adjustments following input loss found in cone-DTR retina demonstrate how functional homeostasis restores neuronal activity to a set point following perturbation. Homeostatic plasticity in the retina has been observed broadly: (1) potentiation of rod-to-rod bipolar cells in the rod degeneration *P23H* mice to maintain light sensitivity (Leinonen et al., 2020), (2) structural and functional compensation at second-order synapses upon partial loss of photoreceptors (Care et al., 2019, 2020; Shen et al., 2020; Lee et al., 2022), (3) preserved cone-mediated light responses and fidelity of information transmission in a *CNGB1* model of rod degeneration (Scalabrino et al., 2022), (4) maintained light responses in cones lacking outer segments in the rod degeneration *rd10* mouse (Xu et al., 2022; Ellis et al., 2023), and (5) retained signal transmission and processing capacity of surviving circuitry in the rod degeneration *rd1* mouse (Rodgers et al., 2023). Despite differences in the etiologies of photoreceptor death, these studies provide evidence that post-receptoral circuits can retain function during degeneration. The present observation of changes in temporal filters following partial cone loss emphasizes that such changes would be missed by conventional diagnostic tools for acuity which tend not to limit decision time for letter identification (Ferris et al., 1982; Ratnam et al., 2013; Foote et al., 2018). Such resiliency within the retina inspires efforts to develop (1) diagnostics sensitive to mild changes and (2) strategies to preserve surviving circuits to restore visual function. For example, computational models of morphologically accurate healthy and degenerated retina revealed maintained responsiveness of ON and OFF cone bipolar cells to electrical stimulation in degenerated retina (Farzad et al., 2023). Based on the resilience of cone vision, cone preservation would be a promising therapeutic strategy. However, further studies are required to determine the time window for effective intervention as a circuit’s rewiring capacity, cone-mediated light responses, and visual behaviors all decline with age (Shen et al., 2020).

The present study demonstrates that while retinal pathways undergo distinct functional and structural modifications, overall the ganglion cells maintain general center-surround receptive field structures and the fidelity of signaling despite input loss, demonstrating compensatory changes that minimally degrade function. Such adaptive changes following input loss have been observed across sensory circuits. For example, in the auditory system, reduced input following noise-induced damage to the synapses of hair cells in the inner ear (cochlear synaptopathy) led to an increase in neural gain across a wide range of frequencies to compensate for input loss (Barbour, 2020; Monaghan et al., 2020). Similarly, acoustic trauma caused auditory cortical neurons to exhibit increased tone-evoked responses as a result of dynamic properties of inhibition, which led to the expansion of receptive fields (Scholl and Wehr, 2008). Another study reported the recovery of functional patterns of auditory cortical activity potentially via changes in the balance of excitatory and inhibitory inputs in adult-onset deafness (Karoui et al., 2023). Across the auditory and visual systems, inhibitory neurons exhibit greater dynamics than excitatory neurons (Villa et al., 2016; Eavri et al., 2018). For example, in the visual system, greater flexibility of inhibitory circuits were found both in the visual cortex and in the retina as reported here. Even beyond pathways within one sensory system, robust plasticity in the form of cortical reorganization has been observed in compensation across sensory modalities in the mature nervous system (Jitsuki et al., 2011; Petrus et al., 2014; Ewall et al., 2021; Kral and Sharma, 2023).

Previous studies have shown that neurons exhibit different vulnerability and plasticity upon deafferentation (O’Brien et al., 2014; Ou et al., 2016; Villa et al., 2016; Eavri et al., 2018; Tran et al., 2019a; Wang et al., 2021; Karoui et al., 2023; Kumar et al., 2023; Dyszkant et al., 2025a), but how each cell type responds and where these changes occur within the circuits have not been examined. By investigating spatiotemporal properties of two types of A_OFF_ ganglion cells, we demonstrate unique outcomes in response to common input loss and provide mechanistic insights on those changes across recording configuration through pharmacology.

## MATERIALS AND METHODS

### Mice

All procedures were done in accordance with the University of California, San Francisco Institutional Animal Care and Use protocols. Animals were maintained on a 12 h dark-12 h light cycle and fed standard mouse diet ad libitum. The following transgenic mouse lines were crossed: *OPN1MWCre* (Le et al., 2004) for Cre-recombinase expression in cones expressing middle wavelength sensitive (M)-opsin, with *Rosa26-loxP-stop-loxP-DTR* (Buch et al., 2005) for Cre-dependent expression of diphtheria toxin receptor (DTR). All transgenic mice were backcrossed into the C57BL/6J background. Male and female mice were used for experiments.

### Diphtheria toxin induced cone death

To induce cone death, one intramuscular diphtheria toxin injection was made between P30-40 at a dosage between 50 ng/g and 75 ng/g for OPN1MW-Cre (Care et al., 2019; Lee et al., 2022). Dosages were adjusted to induce approximately 50% cone loss. Control littermates included either DTR-negative (N = 58) or DTR-positive littermates (N = 11) injected with an equivalent volume of saline. We found no difference in the cone densities between these control groups. Experiments were performed within 55 days after the injection. We did not find significant changes to spatiotemporal properties across the shorter and longer halves of the interval.

### Tissue preparation for immunostaining

Immunostaining protocols were identical to those described previously (Care et al., 2019; Lee et al., 2022). The retinas were fixed in 2% paraformaldehyde for 20 min at room temperature, rinsed in PBS, pH 7.42, then immersed in blocking solution (Jackson Immunoresearch, Cat# NC9624464) overnight, incubated in primary antibodies for 5 days at 4°C, then rinsed in PBS and incubated in secondary antibodies for 1 day at 4°C, and rinsed with PBS and mounted with Vectashield (Vector laboratories Cat# H-1000, RRID: AB_2336789) underneath a coverslip. The primary antibodies utilized in this study were as follows: anti-cone arrestin (Millipore Cat# AB15282, RRID:AB_1163387), anti-gephyrin (Synaptic System Cat# 147-111, RRID: AB_887719), anti-GABA_A_Rβ2,3 (Millipore Cat# MAB341, RRID: AB_2109419), anti-GlyRɑ1 (Synaptic System Cat# 146111: RRID: AB_887723), anti-synaptotagmin II (Zebrafish International resource center Cat# znp-2, RRID: AB_10013783), anti-HCN4 (Alomone Labs Cat# APC-052, RRID: AB_2039906), anti-Lucifer Yellow (Life Technologies Cat# A-5750, RRID: AB_2536190), anti-CtBP2 (BD Bioscience Cat# 612044, RRID: AB_399431), anti-Calbindin (Swant Cat# CB38, RRID: AB_10000340), anti-RBPMS (PhosphoSolutions Cat# 1830-RBPMS, RRID: AB_2492225), anti-PKC (Sigma-Aldrich Cat# P5704, RRID: AB_477375), anti-ChAT (Millipore Cat# AB144P-200UL, RRID: AB_90661), PNA-647 (Life Technologies Cat# L32460). Secondary antibodies used were as follows: anti-mouse-Dylight 405 (Jackson Immunoresearch Cat# 715-475-150, RRID: AB_2340839), anti-mouse-Alexa 647 (Jackson Immunoresearch Cat# 715-605-151, RRID: AB_2340863), anti-rabbit-Alexa 488 (Jackson Immunoresearch Cat# 711-545-152, RRID: AB_2313584).

### Image acquisition

Type 2 and 3a OFF cone bipolar cells and their associated synaptic puncta were imaged on a Leica SP8 confocal microscope using a 40x 1.3 numerical aperture (NA) oil-immersion objective. Image stacks were acquired with a voxel size of 0.09 µm (x axis, y axis) and 0.3 µm (z axis). Each plane was acquired 2 times to obtain the average. Individual dye-filled A_OFF-S_ and A_OFF-T_ ganglion cells were localized using epifluorescence and their dendrites and associated synaptic puncta were imaged with the following settings: 40x (1.3 NA) objective at a resolution of 0.102 x 0.102 x 0.3 µm. A_OFF-S_ and A_OFF-T_ ganglion cells were identified by stratification level of their dendrites (Krieger et al., 2017). To determine how other retinal neurons were affected by the cone loss, we quantified the following cell types in 5 saline-injected DTR-negative and 6 DT-injected DTR-positive mice: horizontal cells, rod bipolar cells, ON starburst amacrine cells, OFF starburst amacrine cells, and ganglion cells (Figure S1E-K). Flat-mounted retina were split along the dorsal-ventral axis to maximize different combinations of antibodies. The following imaging parameters were used for each of the channels: 40x (1.3 NA) objective at a resolution of 0.09 x 0.09 x 0.4 µm. For each channel, we acquired 3 images in different locations for each retina. Each location was selected to be between 1 and 2 mm from the center of the retina, i.e., optic nerve head.

### Electrophysiology tissue preparation

Procedures for patch-clamp recording of A_OFF-S_ and A_OFF-T_ ganglion cells in flat mount retina were identical to those described previously (Care et al., 2019, 2020; Lee et al., 2022). For bar noise stimuli (Figures 2-8), recordings were made in dorsal-nasal retina where the largest A-type ganglion cells reside (Sawant et al., 2021) and where middle (M)-wavelength sensitive opsin dominates (Applebury et al., 2000). Following recordings, retinas were mounted on filter paper, fixed in 2% paraformaldehyde for 15-20 min, and processed for immunostaining.

### Patch-clamp recordings

Patch-clamp recordings from ganglion cells were identical to those described previously (Lee et al., 2022). Patch electrodes were pulled from borosilicate glass (Sutter instruments) on a Narishige puller to 3–6 MOhm resistance. Under infrared light (950 nm), large round somas were targeted as putative A_OFF-S_ ganglion cells and large oblong somas were targeted as putative A_OFF-T_ ganglion cells. To type cells, extracellular spikes were recorded with an electrode filled with HEPES-buffered Ames in cell-attached configuration. A_OFF-S_ ganglion cells were identified by their characteristically sustained spiking before and after a 500 ms light increment. A_OFF-T_ ganglion cells were identified by their characteristically transient spiking after a 500 ms light increment. Once the ganglion cell was identified, a new electrode was used to make an intracellular recording. Whole-cell recordings were made with an electrode filled with internal solution containing either potassium aspartate for current-clamp recordings (Figures 4G-I, 6H-M, S2G-R, S4F) or cesium methane sulfonate for voltage-clamp recordings when acquired in the same cell (Figures 2A-E, J, 3, 4A-C, 5, 6B-G, 8, S2C-F, S4D) and 0.04% Lucifer yellow dye (Care et al., 2019; Lee et al., 2022). Voltage-clamp recordings were made before and after the application of 100µM picrotoxin (Thermo Fischer Scientific P1675-1G) and 5µM strychnine (Sigma-Aldrich S8753) (Figures 3, 5, 8). For perforated patch-clamp recordings in Figure S5, amphotericin-B (0.05 mg/ml) was added to the potassium aspartate internal solution in the back portion, but not the front tip, of the electrode to allow a delay before perforating the cell membrane (Care et al., 2020; Lee et al., 2022).

### Light stimuli

To elicit spike responses for cell identification, a 470 nm LED was used to deliver a uniform spot 500 µm in diameter for 500 ms using Matlab. This wavelength was used to preferentially stimulate rods between the ranges of 8–400 photoisomerizations per rod per sec (Rh*/rod/sec). Procedures for measuring the spatiotemporal receptive field profiles were identical to those described previously (Lee et al., 2022). To preferentially elicit cone-mediated responses, we presented flickering bars (width of 40µm; length of 1000µm) with intensities drawn from a Gaussian white noise distribution with mean intensity of 8,400 Rh*/rod/sec to adapt rods (Care et al., 2019; Lee et al., 2022). For cone-mediated stimuli, a 30ms decrement flash of the 405nm LEDs was presented on a mean background of 4000 rod isomerizations per rod per second (Rh*/rod/sec). The stimuli were presented through a circular aperture 500µm in diameter (Figure 2J).

### Intrinsic excitability of OFF ganglion cells

For current injection experiments (Figure S5B-G), procedures were identical to those described in Lee et al. 2022 (Lee et al., 2022). Spike threshold and resting membrane potential were calculated by injecting a slow ramp of current (Figure S5H-I). Resting membrane potential was taken as the initial membrane voltage 500ms before injecting a linear current injection ramp from 0 to 300pA and spike threshold was the average voltage at which a cell initiated its first action potential in response to the ramp.

### Quantification of cell type numbers

Procedure for counting cones, labeled either by cone arrestin or peanut agglutinin, were identical to those described previously (Care et al., 2019; Lee et al., 2022). Briefly, images (387µm x 387µm) of cones immediately above the recorded ganglion cells were quantified in Imaris. Procedure for counting horizontal cells, starburst amacrine cells (SACs), rod bipolar cell axon stalks, and ganglion cells were the same as for counting cones. Only the horizontal cells were counted manually and averaged across two analyzers because of poor signal to noise with the calbindin staining. Images with especially poor signal to noise ratios were not counted and omitted from the quantified dataset.

### Quantification of synaptic density

To quantify synaptic puncta associated with specific cell types, we used confocal images of T2 and T3a OFF cone bipolar cells labeled by immunostaining for Syt2 and HCN4 in the OFF sublamina, respectively, and individual ganglion cells filled with Lucifer Yellow during physiology experiments. ImageJ (NIH RRID: SCR_003070) was used to median filter the images to remove thermal noise generated by the microscope’s detectors. We created a binary mask of bipolar cell axons of interest (Amira, Thermo-Fisher Scientific RRID: SCR_014305) and VolumeCut (Della Santina et al., 2021) was used to select the OFF sublamina labeling of Syt2 and HCN4 within the IPL. Synaptic puncta associated with bipolar cell axons, A_OFF-S_ and A_OFF-T_ ganglion cell dendrites were identified and manually validated in ObjectFinder (Della Santina et al., 2013) with methods previously used (Lee et al., 2022). Puncta distribution of PSD95, GlyRɑ1, and GABA_A_β2,3 within the dendrites of individual A_OFF-S_ and A_OFF-T_ ganglion cells were identified and quantified using ObjectFinder. Imaris was used to generate a 3-dimensional dendritic skeleton of the ganglion cell. Custom-written Matlab (Mathworks RRID: SCR_001622) routines were used to create a binary mask and used as a criteria to include CtBP2, GlyRɑ1, and GABA_A_β2,3 that exceeds 50% overlap with a mask. Linear puncta density was calculated within a moving window of 10 µm along the dendritic skeleton, starting from the cell soma.

### Linear-nonlinear model

Procedures for analyzing spatiotemporal receptive fields of spikes, subthreshold voltages, excitatory, and inhibitory currents were identical to those described previously (Lee et al., 2022). As in the previous work, we normalized the linear filter such that it was a unit vector using Igor Pro (RRID:SCR_000325). This choice of normalization places all gain information into the nonlinearity. In response to the bar stimulus, the linear filter was used to measure temporal information. A single temporal linear filter was measured by projecting the linear filters from each region of space along the first principal component measured across all filters. A single spatial filter was measured by projecting the linear filters across time along the first principal component across filters. The nonlinear component of the model maps the convolution of the linear filter and stimulus to the cell’s response, i.e., spikes, voltage, excitatory current, or inhibitory current (Figures 6-8).

The linear-nonlinear model was also used in the current injection analysis (Figures S5B-G). The time-reversed spike-triggered average of the injected current represents the linear filter. The nonlinear filter is plotted with the abscissa as the convolution between the spike-triggered average and the stimulus in units of standard deviation against the ordinate, which is the spike rate (Figure S5C). The nonlinearity for each cell was interpolated and smoothed with a spline function before calculating the average.

To evaluate the goodness of fit of the LN model, we calculated the cross correlation coefficients between the response predicted by the model and the data (Kastner and Baccus, 2011). For voltages and currents, continuous responses to the bar noise stimuli were used as is. For spikes, responses were first smoothed with a Gaussian filter with 10ms standard deviation to transform discrete spike times to a continuous response before comparing with the output of the model. Model and data had correlation coefficients for the following recording conditions for each cell type: Control A_OFF-S_ 0.70 ± 0.02 for excitation, 0.66 ± 0.02 for inhibition, 0.38 ± 0.02 for spikes, 0.65 ± 0.02 for voltages; Control A_OFF-T_ 0.67 ± 0.02 for excitation, 0.59 ± 0.02 for inhibition, 0.65 ± 0.02 voltages, 0.36 ± 0.01 for spikes; Cone-DTR A_OFF-S_ 0.66 ± 0.02 for excitation, 0.62 ± 0.02 for inhibition, 0.41 ± 0.02 for spikes, 0.70 ± 0.02 for voltages; Cone-DTR A_OFF-T_ 0.69 ± 0.02 for excitation, 0.61 ± 0.02 for inhibition, 0.41 ± 0.02 for spikes, 0.70 ± 0.02 voltages; Partial A_OFF-S_ 0.69 ± 0.03 for excitation, 0.64 ± 0.03, Control A_OFF-T_ 0.65 ± 0.02 for inhibition, 0.40 ± 0.02 for spikes, 0.70 ± 0.02 voltages; Partial A_OFF-T_ 0.65 ± 0.02 for excitation, 0.62 ± 0.02 for inhibition, 0.38 ± 0.01 for spikes, 0.68 ±0.02 for voltages (mean ± SEM). There was no evidence of any difference in the goodness of fit of the LN model between the following comparisons: control vs. cone-DTR, control vs. partial, and partial vs. cone-DTR (rank sum test). To further validate the LN model, we presented the same repeated noise sequence across conditions and cell types, and evaluated extracellular spikes (Figure S2V).

### Principal components analysis

The temporal filters (Figure 2, S2), spatial filters (Figure 4, S4), or the nonlinearities (Figure 6, S5) were plotted across conditions (i.e., control, cone-DTR, and partial stimulation for A_OFF-S_ and A_OFF-T_ ganglion cells) in n-dimensional space, where n is the number of points in the linear filters or nonlinearities. Before running PCA, the spatial filters were aligned to their centers. We then performed pairwise comparisons of PC1 values between experimental conditions using the Wilcoxon rank-sum test. The following pairwise comparisons were tested separately for A_OFF-S_ and A_OFF-T_ cell types: control vs. partial, control vs. cone-DTR, and partial vs. cone-DTR.

To quantify the magnitude of change within each cell type, we computed the absolute difference in PC1 values between conditions for each cell type, and then calculated the difference between these magnitudes across cell types. For example, |A_OFF-T_(control - partial)| - |A_OFF-S_(control - partial)|, |A_OFF-T_(control - cone-DTR)| - |A_OFF-S_(control - cone-DTR)|, and |A_OFF-T_(partial - cone- DTR)| - |A_OFF-S_(partial - cone-DTR)|. To evaluate the significance of these differences, we bootstrapped null distributions by randomly permuting cell type identities. The resulting distributions were centered on the observed difference of deltas. Histograms were plotted to visualize the null distributions centered at µ, with standard deviation bounds (µ±1σ and µ±2σ) to represent confidence intervals. All p-values from pairwise Wilcoxon tests and difference of delta were corrected for multiple comparisons using the Holm method.

To compare the impact on cone loss nonlinearities of inputs vs. outputs (Figure 7), nonlinearities were plotted across recording configurations for either A_OFF-S_ or A_OFF-T_ ganglion cells. For the pharmacological manipulations, a difference filter was calculated between the temporal (or spatial) filters and nonlinearities before and after the application of the antagonist. Average differences were for display purposes only (Figures 3, 5, 8G, I, L, N). Differences calculated for each individual cell before and after pharmacological manipulations were used for the actual analysis. For both the individual and the difference filters and nonlinearities the dimension of maximum variance was found across conditions, and that vector was assigned the first principal component (PC1). The magnitude of the projection onto the first principal component was compared between conditions: control vs. cone-DTR, control vs. partial, and cone-DTR vs. partial (Figure S3). The results of these comparisons were used in conjunction to assign mechanisms (Figure 1B-E). While we emphasize the comparisons in which significant differences were detectable, some comparisons result in no evidence of differences despite cone loss or partial stimulation, which could indicate recovery from cone loss (Figure 1B). Other combinations of comparisons are interpreted as inconclusive (Figure S2A), for example, spike temporal filters of A_OFF-T_ ganglion cells (Figure S2Q). Such inconclusive results could be consistent with either circuit changes or compensation; however, further studies would be required to distinguish those possibilities.

### Mean difference vector analysis

In the cases where the first principal component was found to be significantly different between control vs. cone-DTR conditions, we parameterized the filters to determine which characteristics contributed most to the observed changes (Figure S2S-U). For temporal filters, we used the following features: time to peak, time to trough, peak amplitude, trough amplitude, and time to zero crossing (Figure 2F-I). For spatial filters, we used the following features: center height, surround height, center width, and surround width (Figure 4D-F, 4J-M). Each filter, whether temporal or spatial, was represented as a point in n-dimensional space, where n is the number of observations (across space or time) within the filter. For each control filter, we then measured the extent to which modifying each feature changed the filter to become more or less like the mean cone DTR filter. This was done by linearly modifying the control filter, and then projecting that modified filter onto the n-dimensional vector that pointed from the mean control filter to the mean cone DTR filter (“mean difference vector”, representing the direction in n-dimensional space that one must travel to go from the average control filter to the average cone DTR filter; Figure S2S-U). For example, if time to peak shifted under cone loss conditions, we created a temporal filter reflecting that degree of change for each control filter, and then projected the positions of these modified filters onto the mean difference vector. The projection values were normalized such that 1.0 corresponded to the position of the mean cone DTR filter and 0.0 corresponded to the position of the mean control filter. Thus, the closer a projection was to 1.0, the more that particular feature modification shifted the filter in the direction of cone DTR filters. This analysis was repeated for each combination of feature modifications, and the mean marginal contribution of each feature was computed across all combinations to reveal the final contribution values. Because not all features were necessarily independent, it was possible to achieve values outside the interval of [0.0, 1.0].

### Statistical Analysis

Data are represented as median ± interquartile range (IQR) in Figures 2C-D, 3G, 3I, 3L, 3N, 4A-B, 4G-H, 5G, 5I, 5L, 5N, 6B-C, 6E-F, 6H-I, 6K-L, 8G, 8I, 8L, 8N, 9B-G, 9I-L, S1D, S2D-E, S2J-K, S2P-Q, S2V, S4D, S4F, S5H-I, as median ± propagated error in Figures 2E, 4C, 4I, 6D, 6G, 6J, 6M, 7, S2F, S2L, S2R, S4E, S4G. A Wilcoxon rank-sum test (abbr. rank sum) with Holm method, for multiple comparisons correction of the p-value, was used to determine significant differences across the three conditions. A permutation test with Holm method, for multiple comparisons correction of the p-value, was used for Figures 2C-E, 3G, 3I, 3L, 3N, 4A-C, 4G-I, 5G, 5I, 5L, 5N, 6B-M, 7, 8G, 8I, 8L, 8N. A two-way ANOVA test was used to determine significant differences between model-predicted and measured responses (Figure S2V). The following asterisks in the figures indicate p values: * ≤ 0.05, ** ≤ 0.01, *** ≤ 0.005.

## AUTHOR CONTRIBUTION

Conceptualization, methodology, software, resources, writing – review & editing, J.Y.L., J.Y., D.B.K, S.C.H, and F.A.D.; formal analysis, J.Y.L., J.Y., D.B,K, S.C.B, L.D.S., J.V.J., and F.A.D.; investigation, visualization, J.Y.L., J.Y., and F.A.D.; data curation; J.Y.L., J.Y., D.B,K.; validation, writing – original draft, supervision, project administration, funding acquisition, J.Y.L. and F.A.D.

## DATA AVAILABILITY

- All data reported in this paper will be shared by the lead contacts upon request.
- Any additional information required to reanalyze the data reported in this paper is available from the lead contact upon request.

## CODE ACCESSIBILITY

- All original code for the image analysis has been deposited at Github and is publically available as of the date of publication. The information is listed in the key resources table.

## SUPPLEMENTAL MATERIAL

Figures S1-S5 and Dataset S1.

## Acknowledgements

We thank Annika Balraj, Jonah Chan, Jeanette Hyer, Yvonne Ou, Fred Rieke, Rachel O.L. Wong for helpful discussions and comments. This work was supported by NIH through the Vision Core Grants P30 EY002162 (UCSF), K99EY033858 (J.Y.L) and EY029772 (F.A.D.) and by foundation grants from McKnight Scholar Award, Research to Prevent Blindness Unrestricted Grant, and All May See.

