## Supplementary Figures for "Partial input loss differentially modifies neural pathways"

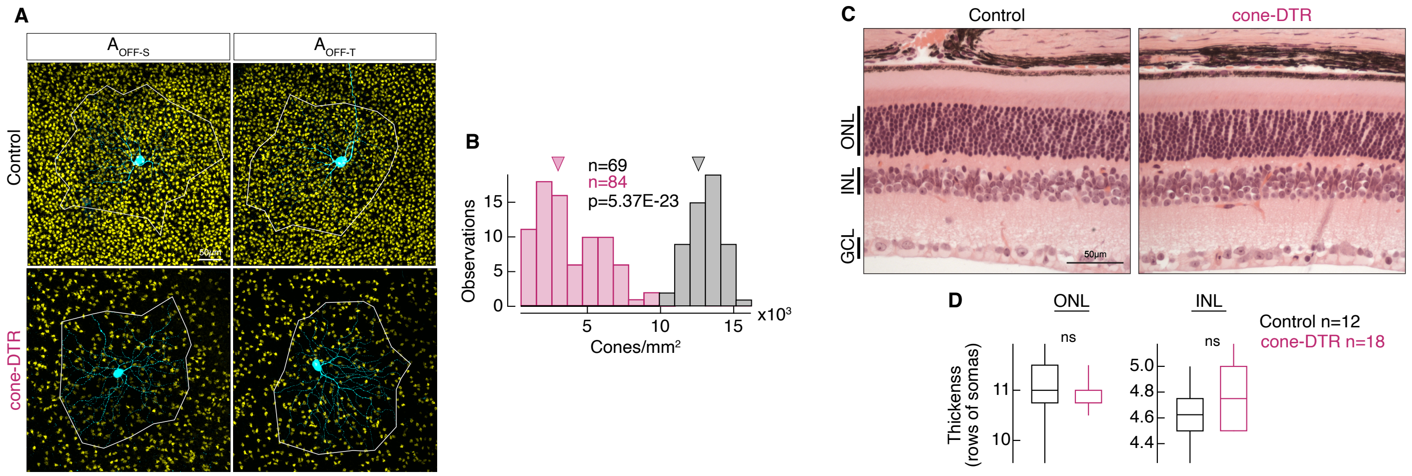


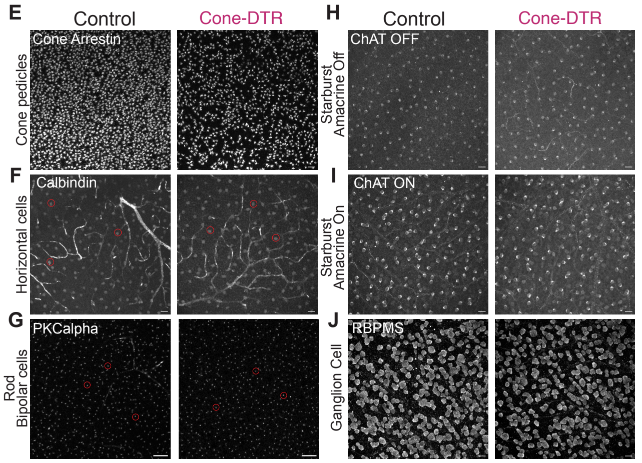


K

| Cell Type | Methods: antibody, retinal layer imaged; retinal topography | Control Density (median ± IQR) | DT-injected Density (median ± IQR) | n = images for control (& DT) N= mice for control (& DT) | Rank sum test p-value |
|---|---|---|---|---|---|
| Cone pedicles | Cone arrestin; OPL; dorsal nasal | 14568.16 ± 1761.50 cells/mm² | 2550.68 ± 4756.71 cells/mm² | n = 12 (18) N = 4 (6) | p=2.40E-06 |
| Horizontal Cells | Calbindin; OPL; dorsal nasal | 919.04±179.81- cells/mm² | 859.11±133.19 cells/mm² | n = 9 (13) N = 3 (5) | p = 0.12 |
| Rod Bipolar Cells | PKCalpha; IPL; dorsal temporal | 14611.45 ± 1351.93 cells/mm² | 14291.78 ± 939.02 cells/mm² | n = 9 (18) N = 3 (6) | p=0.36 |
| Starburst Amacrine Cells (OFF) | ChAT; INL; dorsal nasal | 1115.50 ± 241.42 cells/mm² | 1052.24 ± 301.35 cells/mm² | n = 12 (18) N = 4 (6) | p=0.25 |
| Starburst Amacrine Cells (ON) | ChAT; GCL; dorsal nasal | 1072.22 ± 104.89 cells/mm^2 | 1038.92 ± 153.17 cells/mm² | n = 12 (18) N = 4 (6) | p=0.29 |
| Ganglion Cells | RBPMS; GCL; dorsal temporal | 3629.55 ± 825.81 cells/mm² | 3451.51 ± 1283.66 cells/mm² | n = 8 (16) N = 3 (6) | p=0.38 |

**Figure S1. Simian diphtheria toxin receptor system ablates a partial population of cones in the mouse retina.** (A) Confocal images of cone pedicles labeled with cone arrestin in control and cone-DTR conditions in dorsal-nasal retina of $A_{OFF-S}$ and $A_{OFF-T}$ ganglion cells. The bounding box (white line) shows the region of the dendritic field of the cell. (B) Histogram of cone densities: control 13,007.700±1,581.84 cones/mm2; cone-DTR 4,047.030±280.259 cones/mm2; median±IQR, p=5.37E-23, rank sum. Arrowheads indicate medians. n represents the number of images. Number of animals: control=39, cone-DTR=47. (C) Retinal sections stained with hematoxylin and eosin in control (left) and DT-injected cone-DTR (right) conditions. Select retinal layers are labeled: outer nuclear layer (ONL) containing ~97% rod and ~3% cone cell bodies, inner nuclear layer (INL) containing bipolar and amacrine cell bodies, and the ganglion cell layer (GCL) containing displaced amacrine and ganglion cell bodies. (D) Quantification of the number of cell bodies in each column of the ONL and INL for control (black) and DT-injected cone-DTR (magenta) conditions. Box plots show median with IQR and whiskers from 10% to 90% of the data. n represents the number of slices. Number of animals: control=3, cone-DTR=5. See also Dataset S1. (E-J) Example images of cell types under control and cone-DTR conditions in flat mount retina. (E) Cone pedicles labeled by cone arrestin. (F) Horizontal cell somas labeled by calbindin. (G) Rod bipolar cell axon stalks labeled by PKC alpha. (H) Starburst amacrine cell somas in the inner nuclear layer (INL) labeled by ChAT. (I) Starburst amacrine cell somas in the ganglion cell layer (GCL) labeled by ChAT. (J) Ganglion cell somas labeled by RBPMS. All scale bars = 20$\mu$m. (K) Quantification of cell labels shown in E-J.

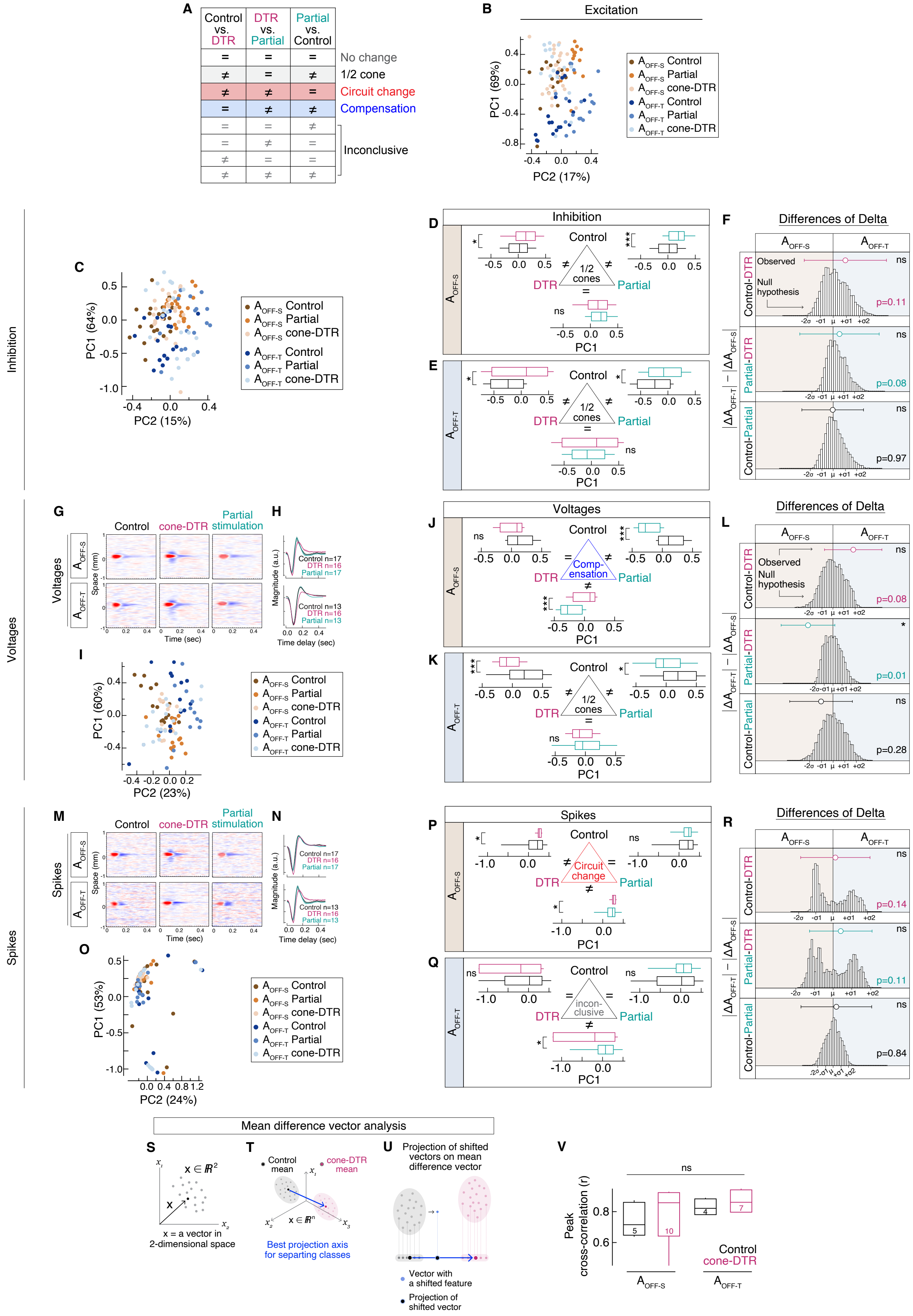
A
Control vs. DTR
DTR vs. Partial
Partial vs. Control
No change
1/2 cone
Circuit change
Compensation
Inconclusive
B
Excitation
PC1 (69%)
PC2 (17%)
AOFF-S Control
AOFF-S Partial
AOFF-S cone-DTR
AOFF-T Control
AOFF-T Partial
AOFF-T cone-DTR
Inhibition
C
PC1 (64%)
PC2 (15%)
D
E
Inhibition
Control
1/2 cones
DTR
Partial
PC1
F
Differences of Delta
Observed
Null hypothesis
Control-DTR
Partial-DTR
Control-Partial
|ΔAOFF-T| − |ΔAOFF-S|
p=0.11
p=0.08
p=0.97
Voltages
G
Control
cone-DTR
Partial stimulation
Space (mm)
Time (sec)
H
Magnitude (a.u.)
Time delay (sec)
Control n=17
DTR n=16
Partial n=17
Control n=13
DTR n=16
Partial n=13
I
PC1 (60%)
PC2 (23%)
J
K
Voltages
Comp-ensation
L
p=0.08
p=0.01
p=0.28
Spikes
M
N
Control n=17
DTR n=16
Partial n=17
Control n=13
DTR n=16
Partial n=13
O
PC1 (53%)
PC2 (24%)
P
Q
Spikes
Circuit change
incon-clusive
R
p=0.14
p=0.11
p=0.84
Mean difference vector analysis
S
x ∈ ℝ²
x = a vector in 2-dimensional space
T
Control mean
cone-DTR mean
x ∈ ℝⁿ
Best projection axis for separting classes
U
Projection of shifted vectors on mean difference vector
Vector with a shifted feature
Projection of shifted vector
V
Peak cross-correlation (r)
Control
cone-DTR
AOFF-S
AOFF-T
ns

**Figure S2. Following partial cone loss, kinetics of temporal filters from inhibitory currents, subthreshold voltages and spikes are similar between $A_{OFF}$ ganglion cell types.** A) Table of possible results in pairwise comparisons of three conditions as the same (=) or different (≠) and the interpretation. Other combinations of comparisons not depicted include additional inconclusive results. (B) Each temporal filter is plotted as a function of its projection onto the first and second principal components from excitation. Variance captured by each principal component noted on the corresponding axis. $A_{OFF\text{-}S}$ plotted in warm colors and $A_{OFF\text{-}T}$ plotted in cool colors for the three conditions. (C) First and second principal components of the inhibitory temporal filters from each $A_{OFF\text{-}S}$ (warm colors) or $A_{OFF\text{-}T}$ (cool colors) across three conditions. Variance captured by each principal component noted on the corresponding axis. (D-E) Box plot of first principal components (PC1) for (D) $A_{OFF\text{-}S}$ and (E) $A_{OFF\text{-}T}$ ganglion cells between conditions. Interpretation of mechanisms in the triangle centers. (F) Difference of deltas results for the null hypothesis for equivalent changes in inhibitory temporal filters between $A_{OFF\text{-}S}$ and $A_{OFF\text{-}T}$ ganglion cells in each comparison of conditions (bootstrapped distribution of 10,000 iterations with noted standard deviations) and the actual difference of deltas (circles with error bars). Significant difference from the null hypothesis displayed on the right. P-values in box plots indicate rank sum comparison between each pair of conditions; p-values are corrected for multiple comparisons with the Holm method (D-E). P-values in the difference of deltas plot from a permutation test with correction for multiple comparisons (F). (G, M) Example spatio-temporal filters in response to a bar noise stimulus under three conditions for either $A_{OFF\text{-}S}$ (top) or $A_{OFF\text{-}T}$ (bottom) under current clamp for measuring (G) voltages and (M) spikes. (H, N) Average temporal filters from (H) voltages (N) spikes of (top) $A_{OFF\text{-}S}$ and (bottom) $A_{OFF\text{-}T}$ ganglion cells in control, cone-DTR, and partial stimulation of control retina. (I, O) First and second principal components of (I) voltage and (O) spike temporal filters from each $A_{OFF\text{-}S}$ (warm colors) or $A_{OFF\text{-}T}$ (cool colors) for the three conditions. Variance captured by each principal component noted on the corresponding axis. (J-K, P-Q) Box plots of first principal components (PC1) for (J, K) $A_{OFF\text{-}S}$ and (P, Q) $A_{OFF\text{-}T}$ ganglion cells between conditions for (I-J) voltages, and (O-P) spikes. Interpretation of mechanisms in triangle centers. (L, R) Difference of deltas results for the null hypothesis for equivalent changes in temporal filters from (L) voltages and (R) spikes between $A_{OFF\text{-}S}$ and $A_{OFF\text{-}T}$ ganglion cells in each comparison of conditions (bootstrapped distribution of 10,000 iterations with noted standard deviations) and the actual difference of deltas (circles with error bars). Significant difference from the null hypothesis displayed on the right. P-values in box plots indicate rank sum comparison between each pair of conditions; p-values are corrected for multiple comparisons with the Holm method (J-K, P-Q). P-values in difference of deltas plots from a permutation test with correction for multiple comparisons (L, R). (S-U) Schematic of temporal filters from each cell plotted in n-dimensional space (S), followed by identifying the mean difference vector between conditions, e.g., control vs. cone-DTR (T), and finally a projection of how shifting the temporal filter according to specific features, e.g., time to peak, time to trough, etc., moves along the vector that discriminates the two conditions of control vs. cone-DTR (U). Peak cross-correlation coefficients (r) between LN model-predicted and measured spike responses to the same repeated noise sequence. Box plots show median with IQR and whiskers from 10% to 90% of the data. See also Dataset S1.

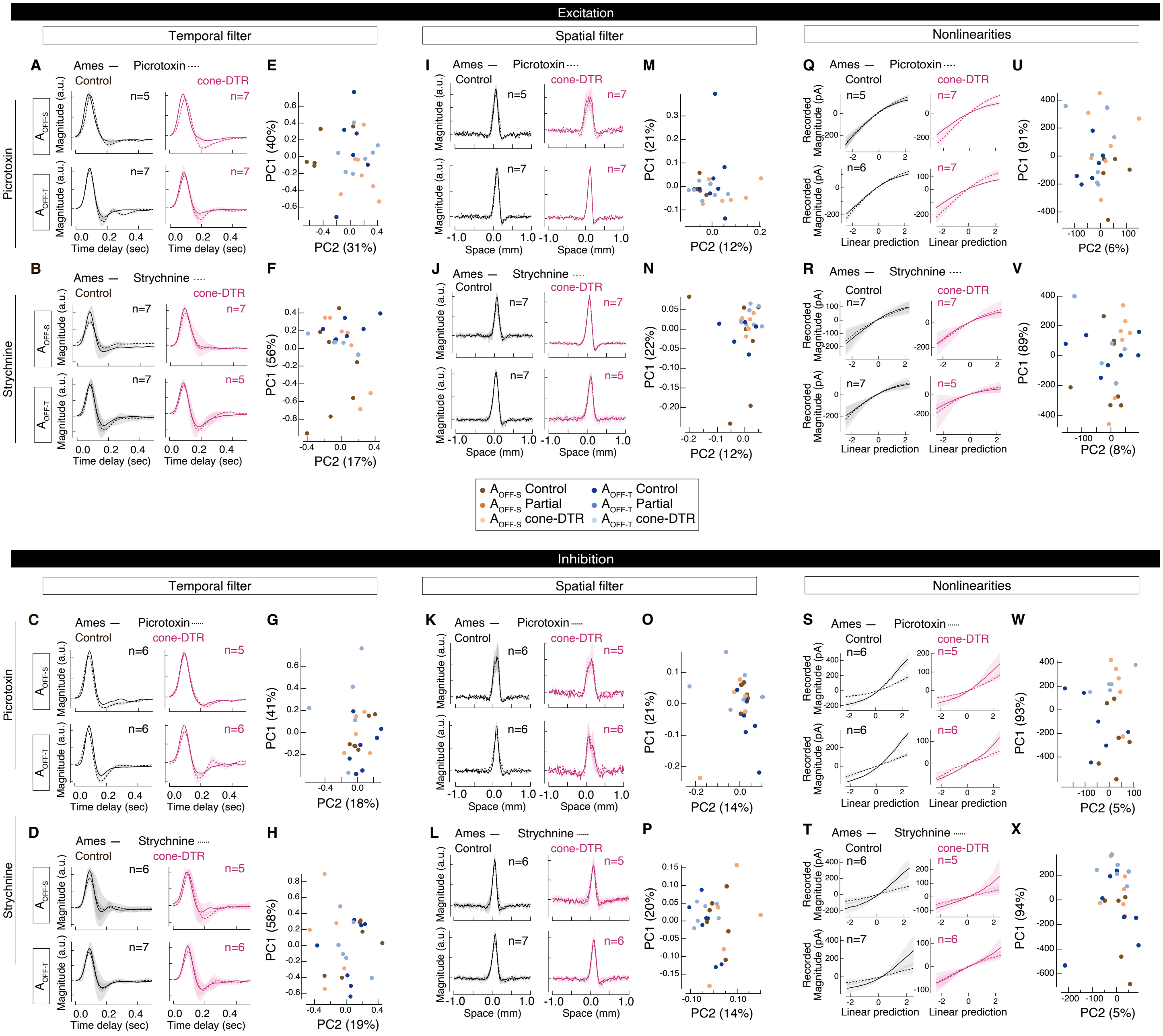


**Figure S3. Principal component analysis on temporal and spatial filters and nonlinearities before and after pharmacological blockade.** (A-D) Average temporal filters from (A-B) excitatory and (C-D) inhibitory currents of $A_{OFF-S}$ (odd rows) and $A_{OFF-T}$ (even rows) ganglion cells in control (left) and cone-DTR (right) retinas under Ames (solid line) and (A, C) 100$\mu$M picrotoxin or (B, D) 5 $\mu$M strychnine (dashed line). (E-H) First and second principal components from (E-F) excitatory and (G-H) inhibitory temporal filter differences between (E, G) picrotoxin and Ames or (F, H) strychnine and Ames of each $A_{OFF-S}$ (warm colors) or $A_{OFF-T}$ (cool colors; refer to box plot colors in the middle). (I-L) Average spatial filters from (I-J) excitatory and (K-L) inhibitory currents of $A_{OFF-S}$ (odd rows) and $A_{OFF-T}$ (even rows) ganglion cells in control (left) and cone-DTR (right) retinas under Ames (solid line) and (I, K) 100$\mu$M picrotoxin or (J, L) 5 $\mu$M strychnine (dashed line). (M-P) First and second principal components from (M-N) excitatory and (O-P) inhibitory spatial filter differences between (M, O) picrotoxin and Ames or (N, P) strychnine and Ames of each $A_{OFF-S}$ (warm colors) or $A_{OFF-T}$ (cool colors). (Q-T) Average nonlinearities from (Q, R) excitatory and (S, T) inhibitory currents of $A_{OFF-S}$ (odd rows) and $A_{OFF-T}$ (even rows) ganglion cells in control (left) and cone-DTR (right) retinas under Ames (solid line) and (Q, S)

100$\mu$M picrotoxin or (R, T) 5 $\mu$M strychnine (dashed line). (U-X) First and second principal components for the (U-V) excitatory and (W-X) inhibitory nonlinearity differences between (U, W) picrotoxin and Ames or (V, X) strychnine and Ames of each $A_{OFF-S}$ (warm colors) or $A_{OFF-T}$ (cool colors). Variance captured by each principal component noted on the corresponding axis. Related to Figures 3, 5, and 8.

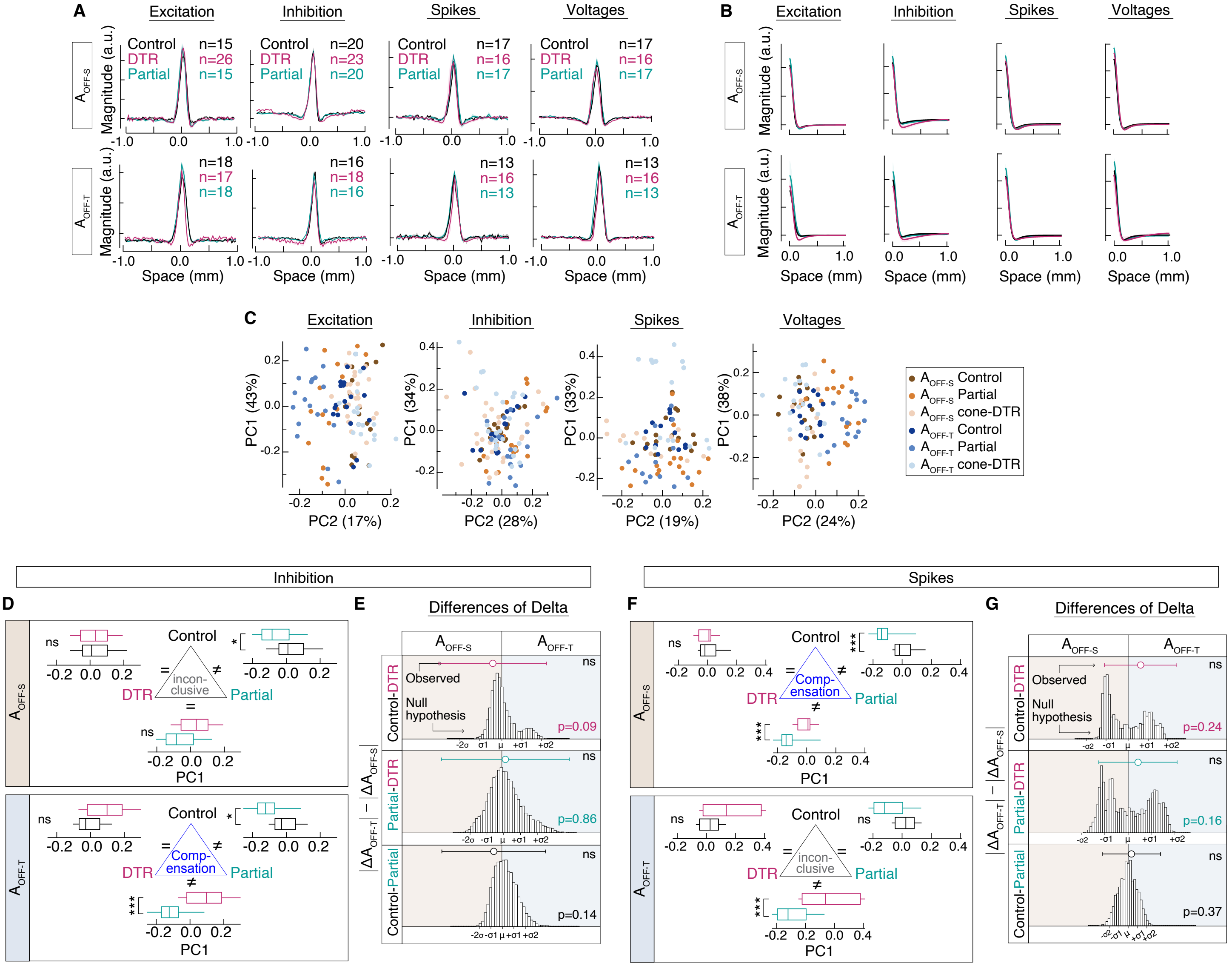


**Figure S4. After partial cone loss, inhibitory and spike spatial filters are similar between $A_{OFF}$ ganglion cell types.** (A, B) Average spatial filters (A) and Difference of Gaussian fit (B) from excitation, inhibition, spikes, and subthreshold voltages of (top) $A_{OFF\text{-}S}$ and (bottom) $A_{OFF\text{-}T}$ ganglion cells in control (black), cone-DTR (magenta) and partial stimulation (green). (C) Projections onto the first and second principal components for the spatial filters of each $A_{OFF\text{-}S}$ (warm colors) or $A_{OFF\text{-}T}$ (cool colors) for the three conditions from excitation, inhibition, spikes, and voltages. Variance captured by each principal component noted on the corresponding axis. (D, F) Box plots of first principal components (PC1) of spatial filters from (D) inhibition and (F) spikes from $A_{OFF\text{-}S}$ (top) and $A_{OFF\text{-}T}$ (bottom) ganglion cells across conditions. Interpretation of mechanisms for excitation in the triangle centers. (E, G) Difference of deltas results for the null hypothesis for equivalent changes in spatial filters from (E) inhibition and (G) spikes between $A_{OFF\text{-}S}$ and $A_{OFF\text{-}T}$ ganglion cells in each comparison of conditions (bootstrapped distribution of 10,000 iterations with noted standard deviations) and the actual difference of deltas (circles with error bars). Significant difference from the null hypothesis displayed on the right. P-values in box plots indicate rank sum comparison between each pair of conditions; p-values are corrected for multiple comparisons with the Holm method (D, F). P-values in difference of deltas plots from a permutation test with correction for multiple comparisons (E, G). See also Dataset S1.

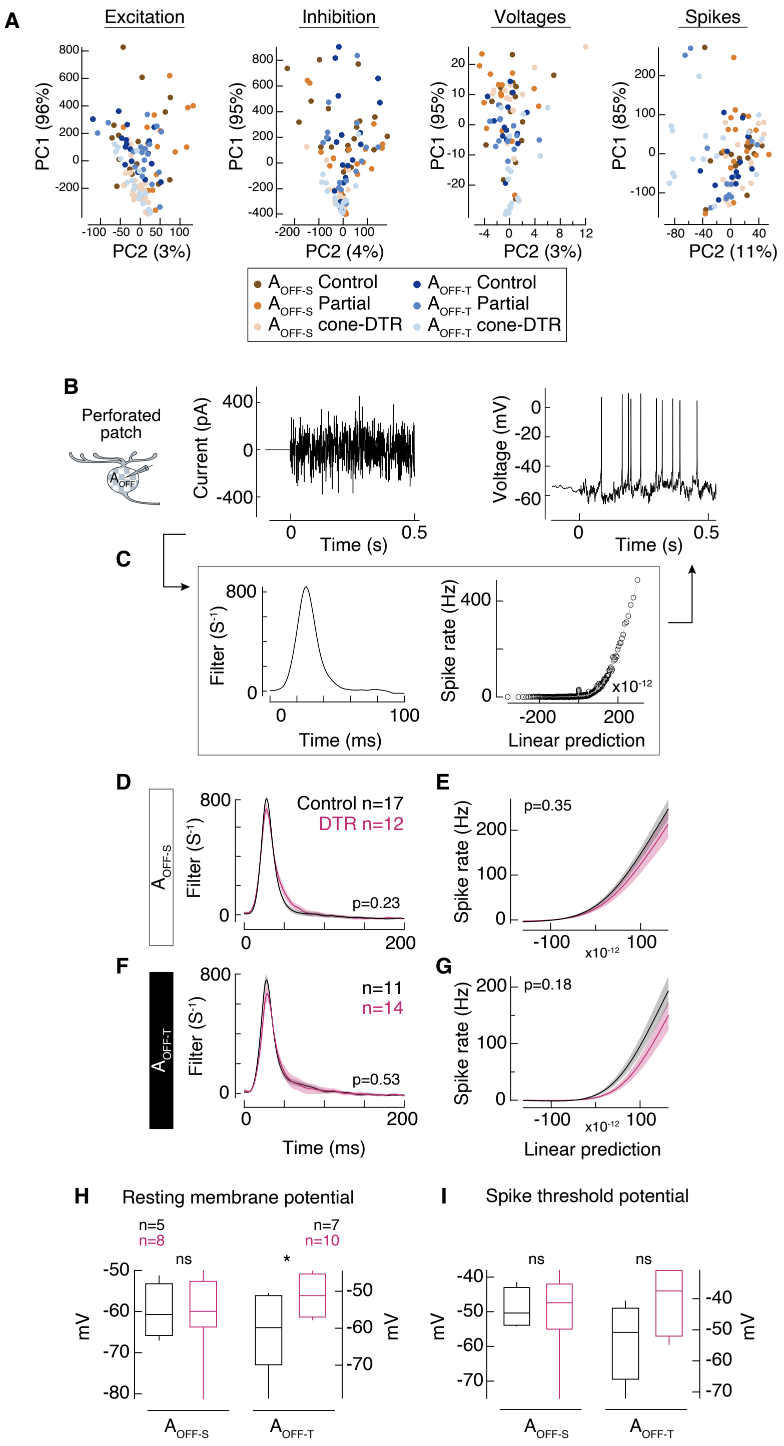


**Figure S5. Intrinsic excitabilities of $A_{OFF-S}$ and $A_{OFF-T}$ ganglion cells are maintained while resting membrane potential of $A_{OFF-T}$ ganglion cells is depolarized after partial cone loss.** (A) First and second principal components for the nonlinearity of each $A_{OFF-S}$ (warm colors) or $A_{OFF-T}$ (cool colors) for the three conditions from excitation, inhibition, voltages, and spikes. Variance captured by each principal component noted on the corresponding axis. (B) $A_{OFF}$ ganglion cells recorded by perforated patch, and current was injected (left) and spikes were recorded (right) on a mean background. (C) Current-to-spike transformation is captured by a time-reversed spike-triggered average (left) and nonlinearity (right), which is fit with a sigmoid function (grey). (D-G) Averages of time-reversed spike-triggered averages and nonlinearities (D-E) of $A_{OFF-S}$ and (F-G) $A_{OFF-T}$

ganglion cells. (H) Resting membrane potential and (I) spike threshold potential were measured from $A_{OFF\text{-}S}$ and $A_{OFF\text{-}T}$ ganglion cells during whole-cell patch-clamp recordings. $A_{OFF\text{-}T}$ ganglion cells of cone-DTR retina were more depolarized than $A_{OFF\text{-}S}$ ganglion cells while spike thresholds were maintained across cell types. Box plots show median with IQR and whiskers from 10% to 90% of the data. n represents the number of cells. See also Dataset S1.
