## Supplementary material for "Partial input loss differentially modifies neural pathways": Dataset S1

| Figure | EXCITATION | | | Full stimulation (Control) | | cone-DTR | | Partial stimulation | | Variance capture by PC1 | Animal number | | |
|---|---|---|---|---|---|---|---|---|---|---|---|---|---|
| | Figure panel | Parameters | | Median | IQR | Median | IQR | Median | IQR | | Control | cone-DTR | Partial stimulation |
| 2 | C | PC1 | OFF-S | 0.092 | 0.256 | 0.172 | 0.471 | 0.346 | 0.272 | 0.687 | OFF-S: 9<br>OFF-T: 10 | OFF-S: 9<br>OFF-T: 8 | OFF-S: 9<br>OFF-T: 10 |
| | D | | OFF-T | -0.512 | 0.404 | 0.225 | 0.345 | -0.246 | 0.334 | | | | |
| | Figure panel | Holm adjusted p-values | | Control vs. cone-DTR | | cone-DTR vs Partial stimulation | | Control vs Partial stimulation | | | | | |
| | C | PC1 | OFF-S | Rank sum | 0.055 | Rank sum | 0.029 | Rank sum | 3.930E-04 | | | | |
| | D | | OFF-T | | 2.436E-06 | | 1.030E-04 | | 0.101 | | | | |
| | Figure panel | Parameters | | Control - cone-DTR | | Partial stimulation - cone-DTR | | Control vs Partial stimulation | | | | | |
| | | | | Median | Propagated Error | Median | Propagated Error | Median | Propagated Error | | | | |
| | E | Difference of Delta | Observed | 0.657 | 0.608 | 0.364 | 0.595 | -0.055 | 0.578 | | | | |
| | | | | Null distribution | | Null distribution | | Null distribution | | | | | |
| | | | Mean | -0.063 | | -3.739E-04 | | -0.033 | | | | | |
| | | | -SD2 | -0.344 | | -0.316 | | -0.313 | | | | | |
| | | | -SD1 | -0.203 | | -0.158 | | -0.173 | | | | | |
| | | | +SD1 | 0.078 | | 0.157 | | 0.107 | | | | | |
| | | | +SD2 | 0.218 | | 0.315 | | 0.247 | | | | | |
| | | | S vs T | Rank sum | 1.330E-04 | Rank sum | 1.250E-05 | Rank sum | 0.746 | | | | |
| | Figure panel | Normalized distance along linear discriminant | | Time to peak | Time to trough | Peak amplitude | Trough amplitude | Time cross zero | | | | | |
| | I | | OFF-S | 0.345 | 0.218 | 0.001 | -0.058 | 0.726 | | | | | |
| | | | OFF-T | 0.132 | 0.210 | 0.026 | 0.208 | 0.830 | | | | | |

| Figure | INHIBITION | | | Control | | cone-DTR | | Partial stimulation | | Variance capture by PC1 | Animal number | | |
|---|---|---|---|---|---|---|---|---|---|---|---|---|---|
| | Figure panel | Parameters | | Median | IQR | Median | IQR | Median | IQR | | Control | cone-DTR | Partial stimulation |
| S2 | D | PC1 | OFF-S | 0.025 | 0.335 | 0.138 | 0.337 | 0.193 | 0.276 | 0.643 | OFF-S: 9<br>OFF-T: 8 | OFF-S: 6<br>OFF-T: 7 | OFF-S: 9<br>OFF-T: 8 |
| | E | | OFF-T | -0.238 | 0.572 | 0.096 | 0.984 | -0.078 | 0.439 | | | | |
| | Figure panel | Holm adjusted p-values | | Control vs. cone-DTR | | cone-DTR vs Partial stimulation | | Control vs Partial stimulation | | | | | |
| | D | PC1 | OFF-S | Rank sum | 0.031 | Rank sum | 0.164 | Rank sum | 0.001 | | | | |
| | E | | OFF-T | | 0.012 | | 0.300 | | 0.015 | | | | |
| | Figure panel | Parameters | | Control - cone-DTR | | Partial stimulation - cone-DTR | | Control vs Partial stimulation | | | | | |
| | | | | Median | Propagated Error | Median | Propagated Error | Median | Propagated Error | | | | |
| | F | Difference of Delta | Observed | 0.221 | 0.742 | 0.119 | 0.743 | -0.008 | 0.574 | | | | |
| | | | | Null distribution | | Null distribution | | Null distribution | | | | | |
| | | | Mean | -0.039 | | 0.017 | | 0.029 | | | | | |
| | | | -SD2 | -0.049 | | -0.333 | | -0.334 | | | | | |
| | | | -SD1 | -0.266 | | -0.158 | | -0.152 | | | | | |
| | | | +SD1 | 0.188 | | 0.192 | | 0.211 | | | | | |
| | | | +SD2 | 0.415 | | 0.367 | | 0.393 | | | | | |
| | | | S vs T | Rank sum | 1.120E-01 | Rank sum | 0.081 | Rank sum | 0.972 | | | | |
| | VOLTAGES | | | Control | | cone-DTR | | Partial stimulation | | Variance capture by PC1 | Animal number | | |
| | Figure panel | Parameters | | Median | IQR | Median | IQR | Median | IQR | | Control | cone-DTR | Partial stimulation |
| | J | PC1 | OFF-S | 0.102 | 0.394 | 0.093 | 0.358 | -0.280 | 0.337 | 0.595 | OFF-S: 8<br>OFF-T: 8 | OFF-S: 3<br>OFF-T: 5 | OFF-S: 8<br>OFF-T: 8 |
| | K | | OFF-T | 0.195 | 0.440 | -0.098 | 0.301 | -0.044 | 0.318 | | | | |
| | Figure panel | Holm adjusted p-values | | Control vs. cone-DTR | | cone-DTR vs Partial stimulation | | Control vs Partial stimulation | | | | | |
| | J | PC1 | OFF-S | Rank sum | 0.044 | Rank sum | 3.900E-04 | Rank sum | 1.420E-05 | | | | |
| | K | | OFF-T | | 0.004 | | 0.589 | | 0.018 | | | | |
| | Figure panel | Parameters | | Control - cone-DTR | | Partial stimulation - cone-DTR | | Control vs Partial stimulation | | | | | |
| | | | | Median | Propagated Error | Median | Propagated Error | Median | Propagated Error | | | | |
| | L | Difference of Delta | Observed | 0.284 | 0.519 | -0.319 | 0.483 | -0.143 | 0.58 | | | | |
| | | | | Null distribution | | Null distribution | | Null distribution | | | | | |
| | | | Mean | -0.082 | | 0.131 | | 0.078 | | | | | |
| | | | -SD2 | -0.481 | | -0.180 | | -0.354 | | | | | |
| | | | -SD1 | -0.282 | | -0.024 | | 0.138 | | | | | |
| | | | +SD1 | 0.118 | | 0.287 | | 0.293 | | | | | |
| | | | +SD2 | 0.318 | | 0.442 | | 0.509 | | | | | |
| | | | S vs T | Rank sum | | Rank sum | | Rank sum | | | | | |
| | SPIKES | | | Control | | cone-DTR | | Partial stimulation | | Variance capture by PC1 | Animal number | | |
| | Figure panel | Parameters | | Median | IQR | Median | IQR | Median | IQR | | Control | cone-DTR | Partial stimulation |
| | P | PC1 | OFF-S | 0.223 | 0.288 | 0.266 | 0.072 | 0.239 | 0.131 | 0.532 | OFF-S: 8<br>OFF-T: 8 | OFF-S: 3<br>OFF-T: 5 | OFF-S: 8<br>OFF-T: 8 |
| | Q | | OFF-T | 0.024 | 0.483 | -0.187 | 1.503 | 0.071 | 0.281 | | | | |
| | Figure panel | Holm adjusted p-values | | Control vs. cone-DTR | | cone-DTR vs Partial stimulation | | Control vs Partial stimulation | | | | | |
| | P | PC1 | OFF-S | Rank sum | 0.039 | Rank sum | 0.017 | Rank sum | 0.236 | | | | |
| | Q | | OFF-T | | 0.051 | | 0.027 | | 0.362 | | | | |
| | Figure panel | Parameters | | Control - cone-DTR | | Partial stimulation - cone-DTR | | Control vs Partial stimulation | | | | | |
| | | | | Median | Propagated Error | Median | Propagated Error | Median | Propagated Error | | | | |
| | R | Difference of Delta | Observed | 0.168 | 1.042 | 0.232 | 0.930 | 0.031 | 0.949 | | | | |
| | | | | Null distribution | | Null distribution | | Null distribution | | | | | |
| | | | Mean | 0.269 | | 0.232 | | 0.049 | | | | | |
| | | | -SD2 | -0.644 | | -0.688 | | -0.252 | | | | | |
| | | | -SD1 | -0.188 | | -0.228 | | -0.102 | | | | | |
| | | | +SD1 | 0.726 | | 0.693 | | 0.199 | | | | | |
| | | | +SD2 | 1.182 | | 0.349 | | 1.153 | | | | | |
| | | | S vs T | Rank sum | 0.144 | Rank sum | 0.844 | Rank sum | 0.109 | | | | |
| | V | Peak cross-correlation between model-predicted and measured spike responses (r) | | Random noise seed | | | | Two-way ANOVA | | | | | |
| | | | | Control | | cone-DTR | | | | | | | |
| | | | | Median | IQR | Median | IQR | conditions 0.705<br>cell types 0.229 | | | | | |
| | | | OFF-S | 0.336 | 0.101 | 0.380 | 0.259 | | | | | | |
| | | | OFF-T | 0.418 | 0.040 | 0.434 | 0.044 | | | | | | |
| | | | | Repeated noise seed | | | | Two-way ANOVA | | | | | |
| | | | | Control | | cone-DTR | | | | | | | |
| | | | | Median | IQR | Median | IQR | conditions 0.473<br>cell types 0.114 | | | | | |
| | | | OFF-S | 0.716 | 0.189 | 0.858 | 0.265 | | | | | | |
| | | | OFF-T | 0.821 | 0.086 | 0.860 | 0.129 | | | | | | |

**Table S1. The first principal components of temporal filters from $A_{OFF\text{-}S}$ and $A_{OFF\text{-}T}$ ganglion cells across conditions. Related to Figures 2, and S2.** (Top) Principal component analysis with Bonferroni-Holm method and mean difference vector analysis on excitatory temporal filters of $A_{OFF\text{-}S}$ and $A_{OFF\text{-}T}$ ganglion cells. Statistical test and results for the comparisons across control, cone-DTR, and partial stimulation. Related to Figure 2. (Bottom) Principal component analysis with Bonferroni-Holm method on inhibitory, spikes and subthreshold voltage temporal filters of $A_{OFF\text{-}S}$ and $A_{OFF\text{-}T}$ ganglion cells. Statistical test and results for the comparisons across control, cone-DTR, and partial stimulation. Related to Figure S2.

| Figure | EXCITATION | | | | | | | Variance capture by PC1 |
|---|---|---|---|---|---|---|---|---|
| | 100uM Picrotoxin | | | Control | | cone-DTR | | |
| | Figure panel | Parameters | | Median | IQR | Median | IQR | |
| 3 | G | PC1 | OFF-S | -0.061 | 0.407 | -0.224 | 0.367 | 0.400 |
| | | | OFF-T | 0.148 | 0.416 | 0.046 | 0.194 | |
| | | Holm adjusted p-values | | Control vs. cone-DTR | | Animal number | | |
| | | | | | | Control | cone-DTR | |
| | | PC1 | OFF-S | Rank sum | 0.114 | OFF-S: 5 | OFF-S: 7 | |
| | | | OFF-T | | 0.365 | OFF-T: 6 | OFF-T: 7 | |
| | 5uM Strychnine | | | Control | | cone-DTR | | Variance capture by PC1 |
| | Figure panel | Parameters | | Median | IQR | Median | IQR | |
| | I | PC1 | OFF-S | -0.155 | 0.794 | 0.178 | 0.504 | 0.561 |
| | | | OFF-T | 0.179 | 0.243 | 0.070 | 0.010 | |
| | | Holm adjusted p-values | | Control vs. cone-DTR | | Animal number | | |
| | | | | | | Control | cone-DTR | |
| | | PC1 | OFF-S | Rank sum | 0.383 | OFF-S: 7 | OFF-S: 7 | |
| | | | OFF-T | | 0.006 | OFF-T: 7 | OFF-T: 5 | |
| | INHIBITION | | | | | | | Variance capture by PC1 |
| | 100uM Picrotoxin | | | Control | | cone-DTR | | |
| | Figure panel | Parameters | | Median | IQR | Median | IQR | |
| | L | PC1 | OFF-S | -0.114 | 0.277 | -0.014 | 0.333 | 0.409 |
| | | | OFF-T | -0.175 | 0.295 | 0.207 | 0.303 | |
| | | Holm adjusted p-values | | Control vs. cone-DTR | | Animal number | | |
| | | | | | | Control | cone-DTR | |
| | | PC1 | OFF-S | Rank sum | 0.931 | OFF-S: 6 | OFF-S: 5 | |
| | | | OFF-T | | 0.014 | OFF-T: 6 | OFF-T: 6 | |
| | 5uM Strychnine | | | Control | | cone-DTR | | Variance capture by PC1 |
| | Figure panel | Parameters | | Median | IQR | Median | IQR | |
| | N | PC1 | OFF-S | 0.101 | 0.674 | 0.197 | 0.566 | 0.584 |
| | | | OFF-T | 1.046E-05 | 0.701 | -0.050 | 0.448 | |
| | | Holm adjusted p-values | | Control vs. cone-DTR | | Animal number | | |
| | | | | | | Control | cone-DTR | |
| | | PC1 | OFF-S | Rank sum | 0.931 | OFF-S: 6 | OFF-S: 5 | |
| | | | OFF-T | | 0.209 | OFF-T: 7 | OFF-T: 6 | |

**Table S2. The first principal components of temporal filters before and after pharmacology from $A_{OFF\text{-}S}$ and $A_{OFF\text{-}T}$ ganglion cells. Related to Figure 3.** Principal component analysis with Bonferroni-Holm method on excitatory and inhibitory current temporal filters of $A_{OFF\text{-}S}$ and $A_{OFF\text{-}T}$ ganglion cells before and after pharmacology. Statistical test and results for the comparisons between control and cone-DTR conditions. Related to Figure 3.

| Figure | EXCITATION | | | Control | | cone-DTR | | Partial stimulation | | Variance capture by PC1 |
|---|---|---|---|---|---|---|---|---|---|---|
| | Figure panel | Parameters | | Median | IQR | Median | IQR | Median | IQR | |
| 4 | A | PC1 | OFF-S | -0.002 | 0.290 | 0.047 | 0.163 | -0.022 | 0.373 | 0.425 |
| | B | | OFF-T | 0.014 | 0.131 | -0.079 | 0.116 | 0.016 | 0.195 | |
| | Figure panel | Holm adjusted p-values | | Control vs. cone-DTR | | cone-DTR vs Partial stimulation | | Control vs Partial stimulation | | |
| | A | PC1 | OFF-S | Rank sum | 0.065 | Rank sum | 0.107 | Rank sum | 0.267 | |
| | B | | OFF-T | | 0.009 | | 0.020 | | 0.938 | |
| | Figure panel | Parameters | | Control - cone-DTR | | Partial stimulation - cone-DTR | | Control vs Partial stimulation | | |
| | | | | Median | Propagated Error | Median | Propagated Error | Median | Propagated Error | |
| | C | Difference of Delta | Observed | 0.044 | 0.264 | 0.026 | 0.308 | -0.017 | 0.333 | |
| | | | | Null distribution | | Null distribution | | Null distribution | | |
| | | | Mean | -0.022 | | -0.023 | | -0.016 | | |
| | | | -SD2 | -0.144 | | -0.179 | | -0.163 | | |
| | | | -SD1 | -0.083 | | -0.101 | | -0.089 | | |
| | | | +SD1 | 0.039 | | 0.055 | | 0.057 | | |
| | | | +SD2 | 0.010 | | 0.132 | | 0.130 | | |
| | | | S vs T | Rank sum | 0.006 | Rank sum | 0.016 | Rank sum | 0.408 | |
| | Figure panel | Normalized distance along linear discriminant | | Center height | Surround height | Center width | Surround width | | | |
| | F | | OFF-T | 0.010 | 0.015 | 0.367 | 0.739 | | | |
| | VOLTAGES | | | Control | | cone-DTR | | Partial stimulation | | Variance capture by PC1 |
| | Figure panel | Parameters | | Median | IQR | Median | IQR | Median | IQR | |
| | G | PC1 | OFF-S | 0.054 | 0.170 | -0.141 | 0.243 | 0.066 | 0.168 | 0.383 |
| | H | | OFF-T | -0.004 | 0.096 | 0.059 | 0.171 | -0.026 | 0.151 | |
| | Figure panel | Holm adjusted p-values | | Control vs. cone-DTR | | cone-DTR vs Partial stimulation | | Control vs Partial stimulation | | |
| | G | PC1 | OFF-S | Rank sum | 0.001 | Rank sum | 0.001 | Rank sum | 0.211 | |
| | H | | OFF-T | | 0.033 | | 0.014 | | 0.325 | |
| | Figure panel | Parameters | | Control - cone-DTR | | Partial stimulation - cone-DTR | | Control vs Partial stimulation | | |
| | | | | Median | Propagated Error | Median | Propagated Error | Median | Propagated Error | |
| | I | Difference of Delta | Observed | -0.131 | 0.239 | -0.121 | 0.258 | 0.010 | 0.207 | |
| | | | | Null distribution | | Null distribution | | Null distribution | | |
| | | | Mean | 0.019 | | 0.008 | | -0.008 | | |
| | | | -SD2 | -0.147 | | -0.169 | | -0.114 | | |
| | | | -SD1 | -0.064 | | -0.080 | | -0.061 | | |
| | | | +SD1 | 0.102 | | 0.097 | | 0.044 | | |
| | | | +SD2 | 0.185 | | 0.186 | | 0.097 | | |
| | | | S vs T | Rank sum | 4.370E-04 | Rank sum | 2.660E-04 | Rank sum | 0.664 | |
| | Figure panel | Normalized distance along linear discriminant | | Center height | Surround height | Center width | Surround width | | | |
| | M | | OFF-S | 0.059 | 0.204 | 0.182 | 0.845 | | | |
| | | | OFF-T | 0.283 | 0.222 | 0.101 | 0.559 | | | |

| Figure | INHIBITION | | | Control | | cone-DTR | | Partial stimulation | | Variance capture by PC1 |
|---|---|---|---|---|---|---|---|---|---|---|
| | Figure panel | Parameters | | Median | IQR | Median | IQR | Median | IQR | |
| S4 | D | PC1 | OFF-S | 0.009 | 0.135 | 0.034 | 0.151 | -0.084 | 0.142 | 0.336 |
| | | | OFF-T | -0.028 | 0.087 | 0.098 | 0.193 | -0.123 | 0.090 | |
| | Figure panel | Holm adjusted p-values | | Control vs. cone-DTR | | cone-DTR vs Partial stimulation | | Control vs Partial stimulation | | |
| | D | PC1 | OFF-S | Rank sum | 0.369 | Rank sum | 0.220 | Rank sum | 0.041 | |
| | | | OFF-T | | 0.059 | | 0.002 | | 0.014 | |
| | Figure panel | Parameters | | Control - cone-DTR | | Partial stimulation - cone-DTR | | Control vs Partial stimulation | | |
| | | | | Median | Propagated Error | Median | Propagated Error | Median | Propagated Error | |
| | E | Difference of Delta | Observed | -0.013 | 0.243 | 0.015 | 0.286 | -0.034 | 0.223 | |
| | | | | Null distribution | | Null distribution | | Null distribution | | |
| | | | Mean | 0.033 | | 0.016 | | 0.015 | | |
| | | | -SD2 | -0.085 | | -0.141 | | -0.087 | | |
| | | | -SD1 | -0.026 | | -0.062 | | -0.036 | | |
| | | | +SD1 | 0.091 | | 0.094 | | 0.065 | | |
| | | | +SD2 | 0.150 | | 0.172 | | 0.116 | | |
| | | | S vs T | Rank sum | 0.092 | Rank sum | 0.143 | Rank sum | 0.856 | |
| | SPIKES | | | Control | | cone-DTR | | Partial stimulation | | Variance capture by PC1 |
| | Figure panel | Parameters | | Median | IQR | Median | IQR | Median | IQR | |
| | F | PC1 | OFF-S | -0.018 | 0.061 | 0.009 | 0.067 | -0.142 | 0.069 | 0.328 |
| | | | OFF-T | 0.026 | 0.123 | 0.138 | 0.397 | -0.119 | 0.141 | |
| | Figure panel | Holm adjusted p-values | | Control vs. cone-DTR | | cone-DTR vs Partial stimulation | | Control vs Partial stimulation | | |
| | F | PC1 | OFF-S | Rank sum | 0.922 | Rank sum | 2.068E-05 | Rank sum | 1.189E-05 | |
| | | | OFF-T | | 0.020 | | 3.575E-05 | | 0.001 | |
| | Figure panel | Parameters | | Control - cone-DTR | | Partial stimulation - cone-DTR | | Control vs Partial stimulation | | |
| | | | | Median | Propagated Error | Median | Propagated Error | Median | Propagated Error | |
| | G | Difference of Delta | Observed | 0.085 | 0.245 | 0.106 | 0.276 | 0.022 | 0.205 | |
| | | | | Null distribution | | Null distribution | | Null distribution | | |
| | | | Mean | 0.068 | | 0.040 | | -0.006 | | |
| | | | -SD2 | -0.207 | | -0.285 | | -0.136 | | |
| | | | -SD1 | -0.069 | | -0.122 | | -0.071 | | |
| | | | +SD1 | 0.206 | | 0.203 | | 0.060 | | |
| | | | +SD2 | 0.344 | | 0.366 | | 0.125 | | |
| | | | S vs T | Rank sum | 2.440E-01 | Rank sum | 0.367 | Rank sum | 0.161 | |

**Table S3. The first principal components of spatial filters from $A_{OFF-S}$ and $A_{OFF-T}$ ganglion cells across conditions. Related to Figures 4 and S4.** (Top) Principal component analysis with Bonferroni-Holm method on excitatory currents and subthreshold voltage spatial filters of $A_{OFF-S}$ and $A_{OFF-T}$ ganglion cells. Statistical test and results for the comparisons across control, cone-DTR, and partial stimulation. Related to Figure 4. (Bottom) Principal component analysis with Bonferroni-Holm method on inhibitory currents and spike spatial filters of $A_{OFF-S}$ and $A_{OFF-T}$ ganglion cells. Statistical test and results for the comparisons across control, cone-DTR, and partial stimulation. Related to Figure S4.

| Figure | | | | | | | | Variance capture by PC1 |
|---|---|---|---|---|---|---|---|---|
| | **EXCITATION** | | | | | | | |
| | 100uM Picrotoxin | | | Control | | cone-DTR | | |
| | Figure panel | Parameters | | Median | IQR | Median | IQR | |
| 5 | G | PC1 | **OFF-S** | -0.012 | 0.007 | -0.048 | 0.069 | 0.213 |
| | | | **OFF-T** | -0.009 | 0.078 | -0.015 | 0.033 | |
| | | Holm adjusted p-values | | Control vs. cone-DTR | | Animal number | | |
| | | | | | | Control | cone-DTR | |
| | | PC1 | **OFF-S** | Rank sum | 0.216 | OFF-S: 5 | OFF-S: 7 | |
| | | | **OFF-T** | | 0.086 | OFF-T: 7 | OFF-T: 7 | |
| | 5uM Strychnine | | | Control | | cone-DTR | | Variance capture by PC1 |
| | Figure panel | Parameters | | Median | IQR | Median | IQR | |
| | I | PC1 | **OFF-S** | 0.008 | 0.169 | 0.022 | 0.025 | 0.221 |
| | | | **OFF-T** | 0.018 | 0.042 | 0.007 | 0.046 | |
| | | Holm adjusted p-values | | Control vs. cone-DTR | | Animal number | | |
| | | | | | | Control | cone-DTR | |
| | | PC1 | **OFF-S** | Rank sum | 0.804 | OFF-S: 7 | OFF-S: 7 | |
| | | | **OFF-T** | | 0.035 | OFF-T: 7 | OFF-T: 5 | |
| | **INHIBITION** | | | | | | | Variance capture by PC1 |
| | 100uM Picrotoxin | | | Control | | cone-DTR | | |
| | Figure panel | Parameters | | Median | IQR | Median | IQR | |
| | L | PC1 | **OFF-S** | 0.024 | 0.053 | 0.025 | 0.063 | 0.209 |
| | | | **OFF-T** | -0.054 | 0.101 | 0.039 | 0.077 | |
| | | Holm adjusted p-values | | Control vs. cone-DTR | | Animal number | | |
| | | | | | | Control | cone-DTR | |
| | | PC1 | **OFF-S** | Rank sum | 0.931 | OFF-S: 6 | OFF-S: 5 | |
| | | | **OFF-T** | | 0.009 | OFF-T: 6 | OFF-T: 6 | |
| | Figure panel | Parameters | | Median | Propagated error | Ranksum | | |
| | 5uM Strychnine | | | Control | | cone-DTR | | Variance capture by PC1 |
| | Figure panel | Parameters | | Median | IQR | Median | IQR | |
| | N | PC1 | **OFF-S** | 0.014 | 0.079 | 0.017 | 0.118 | 0.204 |
| | | | **OFF-T** | 7.000E-03 | 0.090 | 0.025 | 0.047 | |
| | | Holm adjusted p-values | | Control vs. cone-DTR | | Animal number | | |
| | | | | | | Control | cone-DTR | |
| | | PC1 | **OFF-S** | Rank sum | 0.931 | OFF-S: 6 | OFF-S: 5 | |
| | | | **OFF-T** | | 0.122 | OFF-T: 7 | OFF-T: 6 | |

**Table S4. The first principal components of spatial filters before and after pharmacology from $A_{OFF-S}$ and $A_{OFF-T}$ ganglion cells. Related to Figure 5.** Principal component analysis with Bonferroni-Holm method on excitatory and inhibitory current spatial filters of $A_{OFF-S}$ and $A_{OFF-T}$ ganglion cells before and after pharmacology. Statistical test and results for the comparisons between control and cone-DTR conditions. Related to Figure 5.

| Figure | EXCITATION | | | Control | | cone-DTR | | Partial stimulation | | Variance capture by PC1 |
|---|---|---|---|---|---|---|---|---|---|---|
| | Figure panel | Parameters | | Median | IQR | Median | IQR | Median | IQR | |
| 6 | B | PC1 | OFF-S | -140.908 | 386.031 | 197.021 | 120.721 | -127.165 | 307.036 | 0.956 |
| | C | | OFF-T | -111.950 | 185.732 | 183.223 | 269.041 | -65.088 | 266.493 | |
| | Figure panel | Holm adjusted p-values | | Control vs. cone-DTR | | cone-DTR vs Partial stimulation | | Control vs Partial stimulation | | |
| | B | PC1 | OFF-S | Rank sum | 1.267E-06 | Rank sum | 3.911E-05 | Rank sum | 0.935 | |
| | C | | OFF-T | | 1.192E-05 | | 4.450E-04 | | 0.924 | |
| | Figure panel | Parameters | | Control - cone-DTR | | Partial stimulation - cone-DTR | | Control vs Partial stimulation | | |
| | | | | Median | Propagated Error | Median | Propagated Error | Median | Propagated Error | |
| | D | Difference of Delta | Observed | -42.756 | 398.788 | -75.875 | 383.086 | 33.119 | 460.995 | |
| | | | | Null distribution | | Null distribution | | Null distribution | | |
| | | | Mean | 15.450 | | -42.431 | | -91.644 | | |
| | | | -SD2 | -244.946 | | -263.183 | | -281.413 | | |
| | | | -SD1 | -114.748 | | -152.807 | | -186.528 | | |
| | | | +SD1 | 145.647 | | 67.944 | | 3.240 | | |
| | | | +SD2 | 275.845 | | 178.320 | | 98.125 | | |
| | | | S vs T | Rank sum | 0.246 | Rank sum | 0.123 | Rank sum | 0.421 | |
| | INHIBITION | | | Control | | cone-DTR | | Partial stimulation | | Variance capture by PC1 |
| | Figure panel | Parameters | | Median | IQR | Median | IQR | Median | IQR | |
| | E | PC1 | OFF-S | -198.556 | 289.237 | 289.718 | 104.331 | -37.200 | 175.358 | 0.945 |
| | F | | OFF-T | -163.279 | 491.372 | 298.592 | 165.120 | -15.271 | 391.020 | |
| | Figure panel | Holm adjusted p-values | | Control vs. cone-DTR | | cone-DTR vs Partial stimulation | | Control vs Partial stimulation | | |
| | E | PC1 | OFF-S | Rank sum | 4.710E-08 | Rank sum | 6.818E-06 | Rank sum | 0.007 | |
| | F | | OFF-T | | 6.295E-07 | | 2.370E-04 | | 0.045 | |
| | Figure panel | Parameters | | Control - cone-DTR | | Partial stimulation - cone-DTR | | Control vs Partial stimulation | | |
| | | | | Median | Propagated Error | Median | Propagated Error | Median | Propagated Error | |
| | G | Difference of Delta | Observed | -26.402 | 514.169 | -13.054 | 458.470 | -13.347 | 631.600 | |
| | | | | Null distribution | | Null distribution | | Null distribution | | |
| | | | Mean | -28.582 | | -8.619 | | 12.910 | | |
| | | | -SD2 | -292.924 | | -270.177 | | -294.386 | | |
| | | | -SD1 | -160.573 | | -139.398 | | -140.739 | | |
| | | | +SD1 | 103.589 | | 122.160 | | 166.556 | | |
| | | | +SD2 | 235.760 | | 252.940 | | 320.203 | | |
| | | | S vs T | Rank sum | 0.246 | Rank sum | 0.471 | Rank sum | 0.945 | |
| | VOLTAGES | | | Control | | cone-DTR | | Partial stimulation | | Variance capture by PC1 |
| | Figure panel | Parameters | | Median | IQR | Median | IQR | Median | IQR | |
| | H | PC1 | OFF-S | -9.322 | 17.518 | -9.499 | 12.092 | -10.005 | 7.135 | 0.954 |
| | I | | OFF-T | 0.831 | 9.578 | 21.226 | 20.200 | 4.245 | 6.681 | |
| | Figure panel | Holm adjusted p-values | | Control vs. cone-DTR | | cone-DTR vs Partial stimulation | | Control vs Partial stimulation | | |
| | H | PC1 | OFF-S | Rank sum | 0.729 | Rank sum | 0.252 | Rank sum | 0.094 | |
| | I | | OFF-T | | 1.660E-04 | | 0.001 | | 0.031 | |
| | Figure panel | Parameters | | Control - cone-DTR | | Partial stimulation - cone-DTR | | Control vs Partial stimulation | | |
| | | | | Median | Propagated Error | Median | Propagated Error | Median | Propagated Error | |
| | J | Difference of Delta | Observed | 20.220 | 23.161 | 16.475 | 20.927 | 2.732 | 19.899 | |
| | | | | Null distribution | | Null distribution | | Null distribution | | |
| | | | Mean | -2.982 | | -2.272 | | -1.851 | | |
| | | | -SD2 | -16.825 | | -14.233 | | -11.101 | | |
| | | | -SD1 | -9.903 | | -8.253 | | -6.476 | | |
| | | | +SD1 | 3.939 | | 3.708 | | 2.774 | | |
| | | | +SD2 | 10.861 | | 9.689 | | 7.399 | | |
| | | | S vs T | Rank sum | 1.590E-03 | Rank sum | 0.003 | Rank sum | 0.146 | |
| | SPIKES | | | Control | | cone-DTR | | Partial stimulation | | Variance capture by PC1 |
| | Figure panel | Parameters | | Median | IQR | Median | IQR | Median | IQR | |
| | K | PC1 | OFF-S | -0.133 | 38.907 | 27.817 | 118.750 | -29.530 | 65.917 | 0.848 |
| | L | | OFF-T | 43.646 | 66.538 | -16.074 | 108.419 | 77.116 | 133.719 | |
| | Figure panel | Holm adjusted p-values | | Control vs. cone-DTR | | cone-DTR vs Partial stimulation | | Control vs Partial stimulation | | |
| | K | PC1 | OFF-S | Rank sum | 0.211 | Rank sum | 0.012 | Rank sum | 0.091 | |
| | L | | OFF-T | | 0.040 | | 0.015 | | 0.448 | |
| | Figure panel | Parameters | | Control - cone-DTR | | Partial stimulation - cone-DTR | | Control vs Partial stimulation | | |
| | | | | Median | Propagated Error | Median | Propagated Error | Median | Propagated Error | |
| | M | Difference of Delta | Observed | 31.769 | 153.798 | 35.842 | 195.372 | 4.073 | 196.416 | |
| | | | | Null distribution | | Null distribution | | Null distribution | | |
| | | | Mean | -13.371 | | -19.475 | | 6.622 | | |
| | | | -SD2 | -91.104 | | -130.118 | | -67.277 | | |
| | | | -SD1 | -52.238 | | -74.797 | | -30.328 | | |
| | | | +SD1 | 25.496 | | 35.846 | | 43.571 | | |
| | | | +SD2 | 64.362 | | 91.168 | | 80.52 | | |
| | | | S vs T | Rank sum | 7.690E-03 | Rank sum | 0.003 | Rank sum | 0.059 | |

| Figure | OFF-Sustained | | | | | Animal number | | | | | |
|---|---|---|---|---|---|---|---|---|---|---|---|
| | Figure panel | Parameters | | Statistical test | | Control | cone-DTR | | | | |
| | | | | Permutation test | p-value | | | | | | |
| S5 | D | Linear filter | | | 0.232 | 3 | 4 | | | | |
| | E | Nonlinearity | | | 0.350 | | | | | | |
| | OFF-Transient | | | | | Animal number | | | | | |
| | Figure panel | Parameters | | Statistical test | | Control | cone-DTR | | | | |
| | | | | Permutation test | p-value | | | | | | |
| | F | Linear filter | | | 0.532 | 5 | 5 | | | | |
| | G | Nonlinearity | | | 0.178 | | | | | | |
| | Figure panel | Parameters | | Control | | cone-DTR | | Statistical test | | Animal number | |
| | | | | Median | IQR | Median | IQR | Rank sum | p-value | Control | cone-DTR |
| | H | Resting Vm | OFF-S | -60.709 | 9.323 | -59.928 | 9.321 | | 0.840 | 3 | 6 |
| | | | OFF-T | -59.811 | 14.464 | -51.153 | 12.350 | | 0.033 | | |
| | I | Spike threshold Vm | OFF-S | -50.345 | 8.966 | -47.376 | 9.912 | | 0.970 | | |
| | | | OFF-T | -50.824 | 19.964 | -37.466 | 18.107 | | 0.070 | | |

| Figure | Figure panel | Parameters | | Control - cone-DTR | | Partial stimulation - cone-DTR | | Control vs Partial stimulation | |
|---|---|---|---|---|---|---|---|---|---|
| | | | | OFF-Sustained | | | | | |
| | | Parameters | | Median | Propagated Error | Median | Propagated Error | Median | Propagated Error |
| 7 | B | Difference of Delta between Exc and Vol | **Observed** | -337.104 | 332.869 | -322.293 | 301.087 | -13.410 | 402.008 |
| | | | | Null distribution | | Null distribution | | Null distribution | |
| | | | **Mean** | 37.266 | | -31.250 | | -100.856 | |
| | | | **-SD2** | -184.133 | | -201.789 | | -266.861 | |
| | | | **-SD1** | -73.434 | | -116.519 | | -183.858 | |
| | | | **+SD1** | 147.965 | | 54.019 | | -17.853 | |
| | | | **+SD2** | 258.664 | | 139.288 | | 65.15 | |
| | | | **E vs V** | Rank sum | 5.990E-04 | Rank sum | 0.001 | Rank sum | 0.944 |
| | | Parameters | | Control - cone-DTR | | Partial stimulation - cone-DTR | | Control vs Partial stimulation | |
| | | | | Median | Propagated Error | Median | Propagated Error | Median | Propagated Error |
| | | Difference of Delta between Exc and Vol | **Observed** | -275.593 | 222.740 | -229.653 | 239.277 | -45.941 | 227.34 |
| | | | | Null distribution | | Null distribution | | Null distribution | |
| | | | **Mean** | 19.838 | | 12.958 | | -10.176 | |
| | | | **-SD2** | -123.498 | | -132.306 | | -95.681 | |
| | | | **-SD1** | -51.830 | | -59.674 | | -52.929 | |
| | | | **+SD1** | 91.506 | | 85.591 | | 32.576 | |
| | | | **+SD2** | 163.174 | | 158.223 | | 75.328 | |
| | | | **E vs V** | Rank sum | 0.002 | Rank sum | 0.006 | Rank sum | 0.377 |
| | Figure panel | | | OFF-Sustained | | | | | |
| | | Parameters | | Control - cone-DTR | | Partial stimulation - cone-DTR | | Control vs Partial stimulation | |
| | | | | Median | Propagated Error | Median | Propagated Error | Median | Propagated Error |
| | D | Difference of Delta between Exc and Spikes | **Observed** | -317.322 | 319.473 | -296.810 | 288.193 | -8.490 | 385.109 |
| | | | | Null distribution | | Null distribution | | Null distribution | |
| | | | **Mean** | 34.519 | | -25.980 | | -95.395 | |
| | | | **-SD2** | -176.102 | | -188.939 | | -254.235 | |
| | | | **-SD1** | -70.791 | | -107.459 | | -174.815 | |
| | | | **+SD1** | 139.829 | | 55.499 | | -15.976 | |
| | | | **+SD2** | 245.139 | | 136.979 | | 63.444 | |
| | | | **E vs S** | Rank sum | 3.660E-04 | Rank sum | 0.001 | Rank sum | 0.960 |
| | | Parameters | | Control - cone-DTR | | Partial stimulation - cone-DTR | | Control vs Partial stimulation | |
| | | | | Median | Propagated Error | Median | Propagated Error | Median | Propagated Error |
| | | Difference of Delta between Exc and Spikes | **Observed** | -220.874 | 211.144 | -169.147 | 232.680 | -42.132 | |
| | | | | Null distribution | | Null distribution | | Null distribution | |
| | | | **Mean** | 12.226 | | 6.587 | | -5.706 | |
| | | | **-SD2** | -122.589 | | -127.948 | | -87.051 | |
| | | | **-SD1** | -55.181 | | -60.681 | | -46.379 | |
| | | | **+SD1** | 79.634 | | 73.854 | | 34.966 | |
| | | | **+SD2** | 147.041 | | 141.122 | | 75.638 | |
| | | | **E vs S** | Rank sum | 0.004 | Rank sum | 0.018 | Rank sum | 0.308 |
| | Figure panel | | | OFF-Sustained | | | | | |
| | | Parameters | | Control - cone-DTR | | Partial stimulation - cone-DTR | | Control vs Partial stimulation | |
| | | | | Median | Propagated Error | Median | Propagated Error | Median | Propagated Error |
| | F | Difference of Delta between Inh and Vol | **Observed** | -483.820 | 339.451 | -326.621 | 295.433 | -156.369 | 412.822 |
| | | | | Null distribution | | Null distribution | | Null distribution | |
| | | | **Mean** | -13.271 | | -2.365 | | -6.969 | |
| | | | **-SD2** | -166.099 | | -95.706 | | -174.668 | |
| | | | **-SD1** | -89.685 | | -49.036 | | -90.818 | |
| | | | **+SD1** | 63.143 | | 44.305 | | 76.880 | |
| | | | **+SD2** | 139.557 | | 90.976 | | 160.73 | |
| | | | **I vs V** | Rank sum | 3.333E-05 | Rank sum | 9.999E-05 | Rank sum | 0.060 |
| | | Parameters | | Control - cone-DTR | | Partial stimulation - cone-DTR | | Control vs Partial stimulation | |
| | | | | Median | Propagated Error | Median | Propagated Error | Median | Propagated Error |
| | | Difference of Delta between Inh and Vol | **Observed** | -442.717 | 388.189 | -298.667 | 351.925 | -144.050 | 479.357 |
| | | | | Null distribution | | Null distribution | | Null distribution | |
| | | | **Mean** | 9.593 | | 3.876 | | -20.523 | |
| | | | **-SD2** | -213.945 | | -242.593 | | -275.35 | |
| | | | **-SD1** | -102.176 | | -119.359 | | -147.937 | |
| | | | **+SD1** | 121.362 | | 127.111 | | 106.891 | |
| | | | **+SD2** | 233.131 | | 250.346 | | 234.305 | |
| | | | **I vs V** | Rank sum | 2.67E-04 | Rank sum | 0.002 | Rank sum | 0.350 |
| | Figure panel | | | OFF-Sustained | | | | | |
| | | Parameters | | Control - cone-DTR | | Partial stimulation - cone-DTR | | Control vs Partial stimulation | |
| | | | | Median | Propagated Error | Median | Propagated Error | Median | Propagated Error |
| | H | Difference of Delta between Inh and Spikes | **Observed** | -461.790 | 331.729 | -300.587 | 288.474 | -144.762 | 403.027 |
| | | | | Null distribution | | Null distribution | | Null distribution | |
| | | | **Mean** | -13.312 | | -4.331 | | -9.196 | |
| | | | **-SD2** | -159.721 | | -95.867 | | -173.688 | |
| | | | **-SD1** | -86.516 | | -50.009 | | -91.442 | |
| | | | **+SD1** | 59.892 | | 41.347 | | 73.049 | |
| | | | **+SD2** | 133.097 | | 87.024 | | 155.295 | |
| | | | **E vs S** | Rank sum | 9.999E-05 | Rank sum | 3.333E-05 | Rank sum | 0.0815 |
| | | Parameters | | Control - cone-DTR | | Partial stimulation - cone-DTR | | Control vs Partial stimulation | |
| | | | | Median | Propagated Error | Median | Propagated Error | Median | Propagated Error |
| | | Difference of Delta between Inh and Spikes | **Observed** | -410.523 | 382.927 | -256.522 | 350.401 | -140.609 | 471.728 |
| | | | | Null distribution | | Null distribution | | Null distribution | |
| | | | **Mean** | 7.515 | | 2.419 | | -14.787 | |
| | | | **-SD2** | -211.834 | | -240.588 | | -263.405 | |
| | | | **-SD1** | -102.160 | | -110.084 | | -139.096 | |
| | | | **+SD1** | 117.190 | | 123.923 | | 109.522 | |
| | | | **+SD2** | 226.864 | | 245.426 | | 233.831 | |
| | | | **E vs S** | Rank sum | 5.330E-04 | Rank sum | 0.005 | Rank sum | 0.343 |

**Table S5. The first principal components of nonlinearities from $A_{OFF-S}$ and $A_{OFF-T}$ ganglion cells across conditions and linear-nonlinear model for determining the intrinsic excitability of $A_{OFF}$ ganglion cells. Related to Figures 6, S5 and 7.** (Top) Principal component analysis with Bonferroni-Holm method on nonlinearities of $A_{OFF-S}$ and $A_{OFF-T}$ ganglion cells. Statistical test and results for the comparisons across control, cone-DTR, and partial stimulation. Related to Figure 6. (Middle) Statistical tests and results for comparisons of the linear-nonlinear model in response to current injection. Related to Figure S5. (Bottom) Principal component analysis with Bonferroni-Holm method on nonlinearities of input currents and output of either $A_{OFF-S}$ and $A_{OFF-T}$ ganglion cells. Statistical test and results for the comparisons across control, cone-DTR, and partial stimulation. Related to Figure 7.

| Figure | EXCITATION | | | | | | | Variance capture by PC1 |
|---|---|---|---|---|---|---|---|---|
| | 100uM Picrotoxin | | | Control | | cone-DTR | | |
| | Figure panel | Parameters | | Median | IQR | Median | IQR | |
| 8 | G | PC1 | **OFF-S** | -96.373 | 100.429 | -95.679 | 160.493 | 0.908 |
| | | | **OFF-T** | 73.238 | 314.294 | 8.668 | 392.061 | |
| | | Holm adjusted p-values | | Control vs. cone-DTR | | Animal number | | |
| | | | | | | Control | cone-DTR | |
| | | PC1 | **OFF-S** | Rank sum | 0.244 | OFF-S: 5 | OFF-S: 7 | |
| | | | **OFF-T** | | 0.267 | OFF-T: 7 | OFF-T: 7 | |
| | 5uM Strychnine | | | Control | | cone-DTR | | Variance capture by PC1 |
| | Figure panel | Parameters | | Median | IQR | Median | IQR | |
| | I | PC1 | **OFF-S** | -212.396 | 393.590 | 151.980 | 286.304 | 0.898 |
| | | | **OFF-T** | 2.740 | 139.065 | 43.764 | 132.977 | |
| | | Holm adjusted p-values | | Control vs. cone-DTR | | Animal number | | |
| | | | | | | Control | cone-DTR | |
| | | PC1 | **OFF-S** | Rank sum | 0.159 | OFF-S: 7 | OFF-S: 7 | |
| | | | **OFF-T** | | 1.000 | OFF-T: 7 | OFF-T: 5 | |
| | **INHIBITION** | | | | | | | Variance capture by PC1 |
| | 100uM Picrotoxin | | | Control | | cone-DTR | | |
| | Figure panel | Parameters | | Median | IQR | Median | IQR | |
| | L | PC1 | **OFF-S** | -254.479 | 511.076 | 267.947 | 194.059 | 0.932 |
| | | | **OFF-T** | -143.055 | 446.671 | 217.693 | 51.397 | |
| | | Holm adjusted p-values | | Control vs. cone-DTR | | Animal number | | |
| | | | | | | Control | cone-DTR | |
| | | PC1 | **OFF-S** | Rank sum | 0.006 | OFF-S: 6 | OFF-S: 5 | |
| | | | **OFF-T** | | 0.009 | OFF-T: 6 | OFF-T: 6 | |
| | 5uM Strychnine | | | Control | | cone-DTR | | Variance capture by PC1 |
| | Figure panel | Parameters | | Median | IQR | Median | IQR | |
| | N | PC1 | **OFF-S** | -70.773 | 453.683 | 86.927 | 214.795 | 0.935 |
| | | | **OFF-T** | -140.178 | 358.956 | 236.136 | 73.151 | |
| | | Holm adjusted p-values | | Control vs. cone-DTR | | Animal number | | |
| | | | | | | Control | cone-DTR | |
| | | PC1 | **OFF-S** | Rank sum | 0.063 | OFF-S: 6 | OFF-S: 5 | |
| | | | **OFF-T** | | 0.005 | OFF-T: 7 | OFF-T: 6 | |

**Table S6. The first principal components of nonlinearities before and after pharmacology from $A_{OFF\text{-}S}$ and $A_{OFF\text{-}T}$ ganglion cells. Related to Figure 8.** Principal component analysis with Bonferroni-Holm method on excitatory and inhibitory current nonlinearities of $A_{OFF\text{-}S}$ and $A_{OFF\text{-}T}$ ganglion cells before and after pharmacology. Statistical test and results for the comparisons between control and cone-DTR conditions. Related to Figure 8.

| Figure | Figure panel | Parameters | Control | | cone-DTR | | Statistical test | | | |
|---|---|---|---|---|---|---|---|---|---|---|
| | | | Median | IQR | Median | IQR | Statistical test | p-value | | |
| 9 | B | Type 2 CBC volume (µm³) | 11094.000 | 5059.000 | 11418.000 | 4610.500 | Rank sum | 0.488 | | |
| | C | Type 3a CBC volume (µm³) | 7628.000 | 2124.000 | 7859.000 | 2024.000 | | 0.719 | | |
| | Figure panel | Parameters | Control | | cone-DTR | | Statistical test | | Animal number | |
| | | | Median | IQR | Median | IQR | Statistical test | p-value | Control | cone-DTR |
| | D | CtBP2 density, puncta/volume (µm³) | 0.767 | 0.071 | 0.764 | 0.17 | Rank sum | 0.801 | 3 | 5 |
| | | GlyRa1 density, puncta/volume (µm³) | 0.521 | 0.215 | 0.428 | 0.148 | | 0.110 | 4 | 3 |
| | | GABAAβ2 density, puncta/volume (µm³) | 0.660 | 0.218 | 0.631 | 0.087 | | 0.560 | 4 | 4 |
| | E | CtBP2 density, puncta/volume (µm³) | 0.805 | 0.049 | 0.770 | 0.150 | | 0.298 | 3 | 5 |
| | | GlyRα1 density, puncta/volume (µm³) | 0.532 | 0.191 | 0.369 | 0.423 | | 0.029 | 4 | 3 |
| | | GABAAβ2 density, puncta/volume (µm³) | 0.752 | 0.113 | 0.846 | 0.102 | | 0.008 | 6 | 5 |
| | F | CtBP2 volume of puncta (µm³) | 0.150 | 0.060 | 0.200 | 0.050 | Rank sum | 0.003 | | |
| | | GlyRa1 volume of puncta (µm³) | 0.260 | 0.045 | 0.280 | 0.030 | | 0.173 | | |
| | | GABAAβ2 volume of puncta (µm³) | 0.195 | 0.030 | 0.180 | 0.030 | | 0.135 | | |
| | G | CtBP2 volume of puncta (µm³) | 0.190 | 0.020 | 0.200 | 0.070 | | 0.027 | | |
| | | GlyRa1 volume of puncta (µm³) | 0.210 | 0.045 | 0.250 | 0.030 | | 0.089 | | |
| | | GABAAβ2 volume of puncta (µm³) | 0.200 | 0.030 | 0.200 | 0.030 | | 0.581 | | |
| | | Parameters | Control | | cone-DTR | | Statistical test | | | |
| | Figure Panel | Parameters | Median | IQR | Median | IQR | Statistical test | p-value | | |
| | I | Avg PSD95 density by entire IPL (puncta/µm) | 0.732 | 0.115 | 0.808 | 0.066 | Rank sum | 0.455 | | |
| | | Avg Gephyrin density by entire IPL (puncta/µm) | 0.745 | 0.058 | 0.786 | 0.123 | | 0.965 | | |
| | | Avg GlyRα1 density by entire IPL (puncta/µm) | 0.353 | 0.053 | 0.553 | 0.119 | | 0.049 | | |
| | | Avg GABAAβ2,3 density by entire IPL (puncta/µm) | 0.862 | 0.294 | 0.745 | 0.204 | | 0.087 | | |
| | J | Avg PSD95 density by entire IPL (puncta/µm) | 0.888 | 0.069 | 0.742 | 0.044 | | 0.214 | | |
| | | Avg Gephyrin density by entire IPL (puncta/µm) | 0.800 | 0.089 | 0.745 | 0.049 | | 0.233 | | |
| | | Avg GlyRα1 density by entire IPL (puncta/µm) | 0.290 | 0.107 | 0.482 | 0.082 | | 0.022 | | |
| | | Avg GABAAβ2,3 density by entire IPL (puncta/µm) | 0.974 | 0.392 | 0.867 | 0.434 | | 0.974 | | |
| | | Parameters | Control | | cone-DTR | | Statistical test | | Animal number | |
| | Figure Panel | Parameters | Median | IQR | Median | IQR | Statistical test | p-value | Control | cone-DTR |
| | K | PSD95 volume of puncta (µm³) | 0.190 | 0.060 | 0.210 | 0.050 | Rank sum | 0.214 | 15 | 16 |
| | | Gephyrin volume of puncta (µm³) | 0.190 | 0.010 | 0.180 | 0.010 | | 0.199 | 7 | 5 |
| | | GlyRa1 volume of puncta (µm³) | 0.315 | 0.155 | 0.390 | 0.100 | | 0.038 | 6 | 7 |
| | | GABAAβ2 volume of puncta (µm³) | 0.160 | 0.030 | 0.180 | 0.030 | | 0.004 | 7 | 7 |
| | L | PSD95 volume of puncta (µm³) | 0.190 | 0.090 | 0.230 | 0.060 | | 0.017 | 25 | 20 |
| | | Gephyrin volume of puncta (µm³) | 0.185 | 0.025 | 0.160 | 0.030 | | 0.072 | 13 | 7 |
| | | GlyRa1 volume of puncta (µm³) | 0.270 | 0.140 | 0.390 | 0.060 | | 0.021 | 11 | 9 |
| | | GABAAβ2 volume of puncta (µm³) | 0.160 | 0.040 | 0.170 | 0.025 | | 0.015 | 8 | 7 |

**Table S7. Excitatory and inhibitory synapse density on specific bipolar cell types and $A_{OFF}$ ganglion cell types in control and cone-DTR retina. Related to Figure 9.** Statistics of individual type 2 and 3a cone bipolar cell axon volume and excitatory (CtBP2) and inhibitory postsynaptic markers (GlyRα1 and $GABA_AR\beta 2,3$) on type 2 and 3a cone bipolar cell axons. Statistics of individual $A_{OFF\text{-}S}$ and $A_{OFF\text{-}T}$ ganglion cell dendrite-associated postsynaptic excitatory scaffolding protein (PSD95) and inhibitory receptors (GlyRα1 and $GABA_AR\beta 2,3$). Quantification of linear puncta density as a function of dendritic distance from the ganglion cell soma. Statistical tests and results for the comparison between control vs. cone-DTR conditions. Related to Figure 9.
